# Increased PHOSPHO1 expression mediates cortical bone mineral density in renal osteodystrophy

**DOI:** 10.1101/2021.08.06.455411

**Authors:** Shun-Neng Hsu, Louise A Stephen, Scott Dillon, Elspeth Milne, Behzad Javaheri, Andrew A Pitsillides, Amanda Novak, Jose Luis Millán, Vicky E Macrae, Katherine A Staines, Colin Farquharson

**Affiliations:** The Roslin Institute and Royal (Dick) School of Veterinary Studies, University of Edinburgh, Easter Bush, Midlothian, UK; Division of Nephrology, Department of Medicine, Tri-Service General Hospital, National Defense Medical Center, Taipei, Taiwan; Comparative Biomedical Sciences, The Royal Veterinary College, London, UK; Sanford Burnham Prebys Medical Discovery Institute, La Jolla, USA; Centre for Stress and Age-Related Disease, University of Brighton, Brighton, UK

**Keywords:** Bone mineralization, bone mineral density, chronic kidney disease-mineral and bone disorder, renal osteodystrophy, PHOSPHO1, TNAP

## Abstract

Patients with advanced chronic kidney disease (CKD) often present with skeletal abnormalities; a condition known as renal osteodystrophy (ROD). While Tissue-nonspecific alkaline phosphatase (TNAP) and PHOSPHO1 are recognized to be critical for bone mineralization, their role in the etiology of ROD is unclear. To address this, ROD was induced in both wild-type and *Phospho1* knockout (P1KO) mice using dietary adenine supplementation. The mice presented with hyperphosphatemia, hyperparathyroidism, and elevated levels of FGF23 and bone turnover markers. In particular, we noted that in CKD mice, bone mineral density (BMD) was increased in cortical bone (*p* < 0.05) but decreased in trabecular bone (*p* < 0.05). These changes were accompanied by decreased TNAP (*p* < 0.01) and increased PHOSPHO1 (*p* < 0.001) expression in wild-type CKD bones. In P1KO CKD mice, the cortical BMD phenotype was rescued, suggesting that the increased cortical BMD of CKD mice was driven by increased PHOSPHO1 expression. Other structural parameters were also improved in P1KO CKD mice. We further investigated the driver of the mineralization defects, by studying the effects of FGF23, PTH, and phosphate administration on PHOSPHO1 and TNAP expression by primary murine osteoblasts. We found both PHOSPHO1 and TNAP expression to be down-regulated in response to phosphate and PTH. While matrix mineralization was increased with phosphate (Pi), it decreased with PTH and FGF23 had no effect. The *in vitro* data suggest that the TNAP reduction in CKD-MBD is driven by the hyperphosphatemia and/or hyperparathyroidism noted in these mice, while the higher PHOSPHO1 expression may be a compensatory mechanism in an attempt to protect the bone from hypomineralization. We propose that increased PHOSPHO1 expression in ROD may contribute to the disordered skeletal mineralization characteristic of this progressive disorder.

## 1. Introduction

Chronic kidney disease (CKD) is a disorder characterized by progressive loss of kidney function over time. Patients with advanced CKD frequently develop disturbances of mineral and bone metabolism and fail to maintain normal systemic levels of calcium (Ca), inorganic phosphate (Pi), parathyroid hormone (PTH), and fibroblastic growth factor-23 (FGF23) (Moe et al. 2006). Hyperphosphatemia, hyperparathyroidism, and elevated FGF-23 are the primary indicators for the diagnosis of CKD–mineral bone disorder (CKD–MBD) which develops in the early stages of CKD and disease progression can result in cardiovascular disease and renal osteodystrophy (ROD) – the skeletal pathology component of the CKD-MBD syndrome (Fang et al. 2014). The current classification system and treatment strategy for ROD are based on changes to bone turnover, mineralization, and volume (Kazama et al. 2013). A decrease in bone mineral density (BMD) is particularly common in patients with late-stage disease (Nickolas et al. 2013) but animal models have shown a more varied response (Bajwa et al. 2018; Lau et al. 2013a; Metzger et al. 2021). The other ROD-associated skeletal pathologies, have been attributed to CKD-related metabolic and hormonal disturbances (Zheng et al. 2016).

Although the precise mechanisms responsible for the impaired skeletal mineralization observed in ROD remain unclear, the origins are likely to involve a complex interplay between bone and the altered endocrine milieu. Phosphorus retention, due to the failing kidney, leads to chronically elevated concentrations of circulating FGF23 in an attempt to normalize serum Pi levels through enhanced urinary secretion and decreased intestinal absorption (Mirza et al. 2009). This is achieved by the inhibition of renal 1α-hydroxylase and stimulation of 24-hydroxylase but the resulting reduction in circulating levels of 1,25(OH)_2_D_3_ contributes to hypocalcemia and secondary hyperparathyroidism (SHPT) (Shimada et al. 2004). SHPT promotes bone resorption by increasing the receptor activator of nuclear factor-□B ligand (RANKL)/osteoprotegerin (OPG) ratio (Ma et al. 2001). The bone formed during rapid remodeling is both immature and poorly mineralized (Graciolli et al. 2017). Indeed, the mineralization status may be dependent on the prevailing serum PTH concentrations which could explain the various mineralization states reported in ROD (Lau et al. 2013b; Miller et al. 1998). It is also possible that altered endocrine factors may directly target the expression of key phosphatases critical for skeletal mineralization. Specifically, FGF23 may inhibit matrix mineralization by suppressing TNAP expression and activity by osteoblasts resulting in the accumulation of the mineralization inhibitor, pyrophosphate (PPi) (Murali et al. 2016b). Also, PTH may induce a rapid downregulation of *Phospho1* gene expression in osteogenic cells and bone marrow stromal cell lines (Chande and Bergwitz 2018; Houston et al. 2016). Despite clear links between both TNAP and PHOSPHO1 in the control of skeletal mineralization, their roles in ROD remain unclear.

PHOSPHO1 and TNAP are two of the most widely studied phosphatases involved in skeletal mineralization (Dillon et al. 2019). PHOSPHO1 is expressed at sites of mineralization and liberates Pi from phospholipid substrates for incorporation into the mineral phase (Roberts et al. 2007). *Phospho1* deficient mice exhibit decreased BMD, compromised trabecular and cortical bone microarchitecture, and spontaneous greenstick fractures (Boyde et al. 2017). TNAP is an ectoenzyme and hydrolyzes PPi to allow the propagation of hydroxyapatite in the extracellular matrix (ECM), beyond the confines of the matrix vesicle membrane (Hessle et al. 2002). Mice deficient in TNAP (*Alpl*^*-/-*^) phenocopy infantile hypophosphatasia (HPP), an inborn error of metabolism resulting in rickets and osteomalacia (Whyte 2008). A complete absence of ECM mineralization is observed in *Phospho1*^*-/-*^; *Alpl*^*-/-*^ double knockout mice and in murine metatarsals cultured in the presence of PHOSPHO1 and TNAP inhibitors demonstrating the functional co-operativity of PHOSPHO1 and TNAP for bone mineralization (Huesa et al. 2015; Yadav et al. 2011).

Despite great advances in understanding the mechanisms responsible for the altered mineralization status noted in ROD, the involvement of phosphatases is unclear. Therefore, in this study, we examined changes in the expression of PHOSPHO1 and TNAP and bone architecture in long bones using the well-established adenine-induced model of CKD (Jia et al. 2013). We also examined the effects of PTH, FGF23, and Pi on TNAP and PHOSPHO1 expression in primary osteoblasts. Our findings support a specific role for PHOSPHO1particualarly in the altered cortical bone mineralization status in ROD.

## 2. Materials and methods

All reagents were from Sigma-Aldrich (Dorset, UK) or less otherwise stated.

### 2.1. Mice

C57BL/6 male mice (Charles River Laboratories, Currie, UK) and were used in the first *in vivo* study. Male *Phospho1* knockout (P1KO) mice and wild-type (WT) controls, maintained on a C57BL/6 background were generated and genotyped as previously described (Yadav et al. 2011) and used in the second *in vivo* study. At 8-weeks of age, mice were randomly assigned a control (n = 12) or CKD (n = 12) diet (Fig. S1A). Mice losing more than 30% of their body weight were euthanized by exposure to CO_2_ and confirmed dead by cervical dislocation. All animal experiments were approved by the Roslin Institute’s named veterinary surgeon and named animal care and welfare officer (NACWO), with animals maintained in accordance with the Home Office code of practice (for the housing and care of animals bred, supplied, ARRIVE guidelines or used for scientific purposes).

### 2.2. CKD diet and tissue collection

CKD was induced by feeding a casein-based diet containing 0.6% calcium, 0.9% phosphate, and 0.2% adenine (Envigo, Teklad Co. Ltd). Control mice received the same diet without adenine. At 13 weeks of age, all animals were sacrificed, and blood was obtained by cardiac puncture under terminal anesthesia. Femora, tibiae, and kidneys were harvested and processed and stored accordingly.

### 2.3. Serum and urine biochemistry

Serum blood urea nitrogen (BUN), creatinine (Cr), Ca, Pi, and alkaline phosphatase (ALP) activity were quantified using a biochemistry analyzer (Beckman Coulter AU480, Olympus). Intact PTH (Pathway Diagnostics, Dorking, UK), FGF23 (Kainos Laboratories, Inc. Japan), N-terminal propeptide of human procollagen type I (P1NP) and carboxy-terminal telopeptide of type I collagen (αCTX) (Wuhan Fine Biotech, Wuhan, China), levels were determined by ELISA according to manufacturers’ instructions. Hydrophobic bedding, LabSand (Coastline Global, CA, USA) was used to collect urine samples from which. the concentration of Cr and albumin were determined by semi-quantitative test strips (Microalbustix, Siemens) and the specific gravity (SG) was determined by a manual refractometer.

### 2.4. Histopathological analysis of kidney and bone tissues

The right tibiae and kidneys were fixed in 4% paraformaldehyde (PFA, for 24 hrs) and stored in 70% ethanol. Kidneys were processed to paraffin wax using standard procedures. Hematoxylin and eosin (H&E), Masson’s trichrome, and von Kossa staining were performed according to standard methods. Histopathological scoring of renal interstitial inflammation, tubular atrophy, protein casts, and renal fibrosis was defined as 0 = normal; 1 = mild, involvement of < 25% of the cortex; 2 = moderate, involvement of 25 to 50% of the cortex; 3 = severe, involvement of 50 to 75% of the cortex; 4 = extensive, involvement of > 75% of the cortex. Bones were decalcified in 10% ethylenediaminetetraacetic acid (EDTA; pH 7.4) for 14 days at 4°C and processed to paraffin wax. Sections were stained by Goldner’s Trichrome and reacted for tartrate-resistant acid phosphatase (TRAP). Bone histomorphometry was quantified using the BioQuant Osteo software (BIOQUANT Image Analysis Corporation, Texas, USA) using the approved ASBMR histomorphometry nomenclature (three sections/bone: six randomly selected bones from each group).

### 2.5. Microcomputed tomography (μCT)

The bone structure of the left tibiae was determined using micro-computed tomography (μCT, Skyscan 1172, Bruker, Kontich, Belgium). High-resolution scans with an isotropic voxel size of 5 μm were acquired (60 kV, 167 μA, and 0.5 mm filter, 0.6° rotation angle) and from the reconstructed images (NRecon 1.7.3.0 program; Bruker), CTAn software 1.15.4.0 (Skyscan) was used to visualize and determine bone histomorphometric parameters. Three-dimensional images were created using IMARIS 9.0.

In the proximal tibial metaphysis the volume of interest (VOI) extended distally 5% from the bottom of the growth plate excluding the cortical shell. A total of 250 slices beneath this 5% were selected to exclude the primary spongiosa. In the first *in vivo* study, whole bone cortical analysis was performed on datasets derived from whole μCT scans using BoneJ (version 1.13.14) a plugin for ImageJ. Following segmentation, alignment, and removal of fibula from the dataset, a minimum bone threshold was selected for each bone to separate higher density bone from soft tissues and air. The most proximal and the most distal 10% portions of tibial length were excluded from analysis, as these regions include trabecular bone. In the 2^nd^ *in vivo* study cortical analysis was performed on datasets derived from μCT scan images at 50% of the total tibial length from the top of the tibia. BMD phantoms of known calcium hydroxyapatite mineral densities of 0.25 and 0.75 g/cm^3^ were scanned and reconstructed using the same parameters as used for bone samples.

### 2.6. Primary calvarial osteoblast isolation and culture

Calvarial osteoblasts were obtained from 3 to 5-day-old C57BL/6 mice by sequential enzyme digestion [1 mg/ml collagenase type II (Worthington Biochemical, Lakewood, NJ, USA) in Hanks’ balanced salt solution (Life Technologies, Paisley, UK); 4 mM EDTA]. The cells were grown in α-minimum essential medium (αMEM, Invitrogen, Paisley, UK) supplemented with 10% fetal bovine serum (FBS) and 0.5% gentamycin (Life Technologies) until confluent.

### 2.7. Establishment of Pi-substrate free mineralization model for primary osteoblast culture

To study the effects of varying Pi concentrations on phosphatase expression it was essential to control Pi concentration in the basal mineralizing medium. This ruled out the use of β-glycerophosphate (βGP) as the availability of Pi from βGP requires the action of TNAP (Huesa et al. 2015) which can itself be modulated by CKD-associated endocrine factors such as Pi, PTH, and FGF23 (Houston et al. 2016; Rendenbach et al. 2014; Shalhoub et al. 2011). Therefore, upon confluence (day 0), mineralization was induced by supplementing the growth medium (1.8 mM Ca;1 mM Pi) with 50 μg/ml L-ascorbic acid (AA) and 1.5 mM CaCl_2_ (Houston et al. 2016). Cultures were also supplemented with a range of Pi (1-5 mM), PTH (0-50 nM), and FGF23 (0-200 ng/ml) with or without klotho (50 ng/ml) (R&D Systems, Abington, UK). Cells were maintained in a 5% CO_2_ atmosphere at 37°C and mineralization media was changed every second/third day for 28 days.

### 2.8. Cell viability and cytotoxicity assay

To assess the effects of Pi on osteoblast viability, the AlamarBlue assay (Thermo Fisher Scientific, Loughborough, UK), and lactate dehydrogenase (LDH) CytoTox 96 cytotoxicity assay (Promega, Southampton, UK) were performed according to manufacturer’s instructions.

### 2.9. RNA extraction and quantitative polymerase chain reaction

The distal and proximal epiphyses of the left femorae were excised, and the diaphyseal bone marrow was removed by centrifugation at 13,000 x g for 10 mins at 4°C. The resultant cortical shafts were homogenized using a Rotor-Stator Homogenizer (Ultra-Turrax T10). RNA extraction from the homogenized bone and cultured osteoblasts was completed using the RNeasy kit (Qiagen). The RNA concentration was determined using a NanoDrop spectrophotometer (Fisher Scientific, Loughborough, UK) at a wavelength of 260 nm, and RNA purity was evaluated by the 260/280 nm ratio. RNA was reverse transcribed to complementary DNA (cDNA) using Superscript II (Invitrogen). All genes were analyzed with the SYBR green detection method (PCR Biosystems, UK) using the Stratagene Mx3000P real[time QPCR system (Agilent Technologies, Santa Clara, USA). Gene expression data were normalized against housekeeping genes (*Gapdh* in primary osteoblasts and *Atp5b* in bone tissue) using MxPro software (Cheshire, UK). The relative expression of the analyzed genes was calculated and expressed as a fold change compared to control values. Primer sequences are listed in Supplementary Table S1.

### 2.10. Protein extraction and isolation from brush border membrane vesicles (BBMV) of kidney

Kidneys were homogenized in ice-cold buffer A [50 mM D-mannitol, 2 mM 4-(2-hydroxyethyl)-1-piperazineethanesulfonic acid (HEPES), 2.5 mM ethylene glycol-bis (2-aminoethylether)-N, N, N’, N’-tetraacetic acid (EGTA), and 12 mM Tris-base titrated to pH 7.1] mixed with a protease inhibitor cocktail. BBMVs were isolated from microvilli of kidneys using 2 consecutive magnesium precipitations in buffer A and then buffer B [150 mM D-mannitol, 2.5 mM EGTA, and 6 mM Tris hydrochloride. The resultant BBMV pellet was resuspended in radioimmunoprecipitation assay (RIPA) buffer (Thermo Fisher Scientific) containing a protease inhibitor cocktail.

### 2.11. Western blot analysis

Protein from cultured osteoblasts and right femorae diaphyseal cortical bone (with marrow removed) was extracted in RIPA buffer containing protease inhibitor cocktail after homogenization. Protein concentrations were determined using the BCA protein assay kit (Life Technologies). Proteins were separated using a 10% Bis-Tris protein gel (Thermo Fisher Scientific). After blocking in 5% skimmed milk/Tris-buffered saline with Tween 20 (TBST) or LI-COR buffer at room temperature (RT) for 1 hour, the membranes were incubated sequentially with primary and secondary antibodies (Tables S2 and S3). Western blot analysis of proteins from primary osteoblasts was performed using the Odyssey infrared detection system (LI-COR). Western blot analysis of proteins from bone tissues was undertaken using the ultra-sensitive enhanced chemiluminescence detection system (Thermo Fisher Scientific). The blots were imaged by the GeneGnome XRQ chemiluminescence imaging system (Syngene, UK). Densitometry of the protein bands was analyzed with Image J software (NIH) for quantification.

### 2.12. Quantification of ECM mineralization

Cultured osteoblasts were fixed in 4% PFA for 10 mins at RT and stained with aqueous 2% (w/v) Alizarin red solution for 10 mins at RT. The bound stain was solubilized in 10% cetylpyridinium chloride and the optical density was measured by spectrophotometry at 570 nm.

### 2.13. Statistical analysis

Quantitative data are expressed as the mean ± standard error of the mean (SEM) of at least three biological replicates per experiment. The precise number (n) is indicated in the relevant table and figure legends. Statistical analysis was performed using a two-tailed Student’s t-test or one-way analysis of variance (ANOVA) followed by Tukey’s range test, as appropriate. Statistical analysis was implemented by the GraphPad Prism software. A *p* < 0.05 was considered to be significant and noted as “*”. *p* values of < 0.01, < 0.001, and < 0.0001 were noted as “**”, “***” and “****” respectively.

## 3. Results

### 3.1. Verification of the CKD-MBD mouse model

Before investigating TNAP and PHOSPHO1 expression in experimental ROD we first confirmed that our mouse model presents with the characteristic serum biochemistries and kidney pathology of CKD-MBD. The CKD mice lost bodyweight and presented with the expected changes to serum and urine analyte levels at the end of the study (Fig. S1B and Table 1). The kidneys of CKD mice presented with various pathologies including tubular atrophy, protein casts, interstitial inflammation, and renal fibrosis (Fig S2). Furthermore, transcripts encoding kidney injury biomarkers *Lcn2* (protein; Ngal), and *Spp1* [protein; osteopontin (OPN)] (Kaleta 2019; Viau et al. 2010), as well as *Fgf23*, were increased in CKD mice whereas *Slc34a1* (protein; NaPi-2a) expression was decreased (Fig. S3A). Protein expression of OPN and NaPi-2a by BBMV confirmed the transcript data (Fig. S3B). Collectively, these data confirm previous reports that mice fed an adenine-rich diet for 5 weeks developed CKD-MBD (Jia et al. 2013; Metzger et al. 2020; Tamura et al. 2009).

**Table 1.**
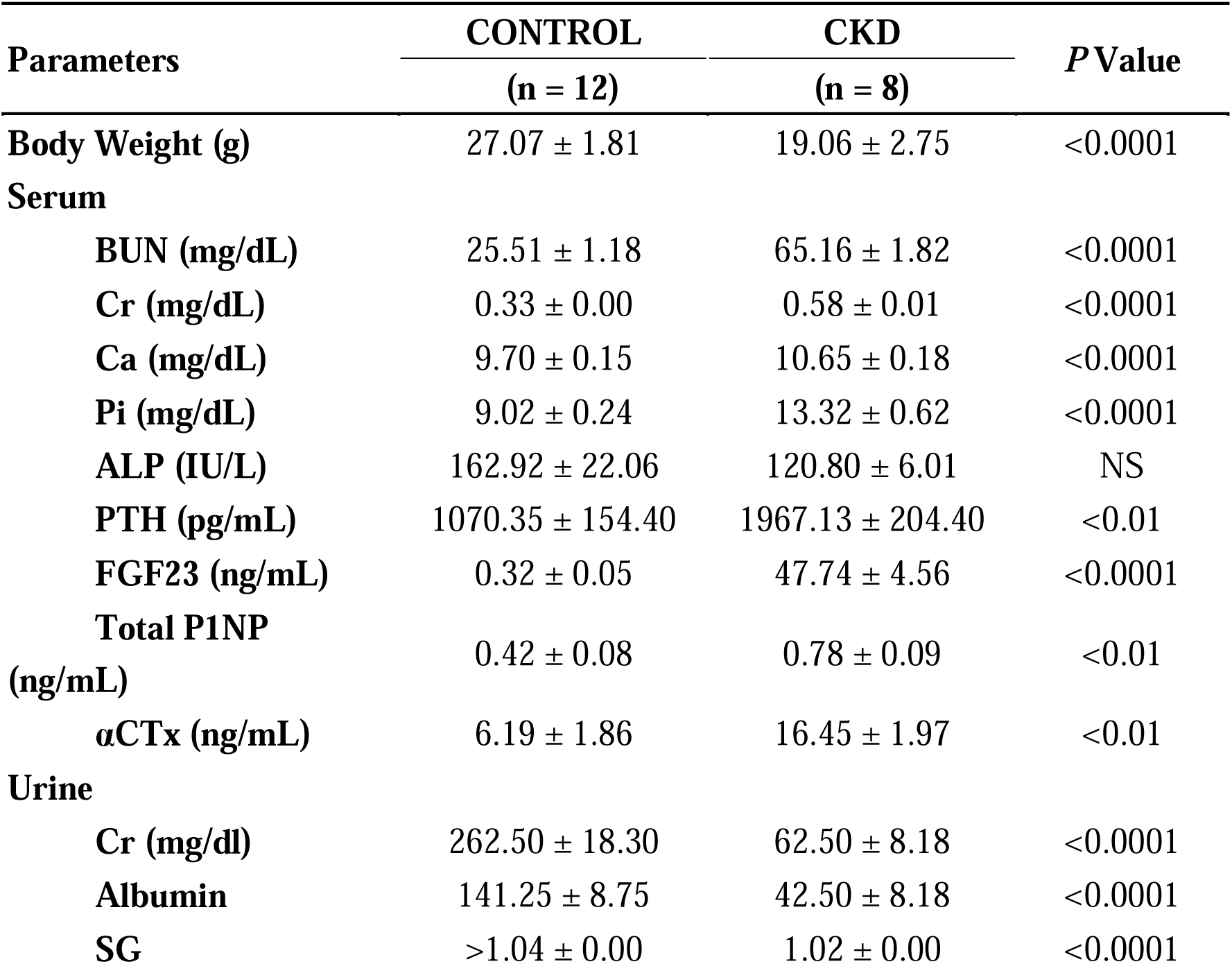
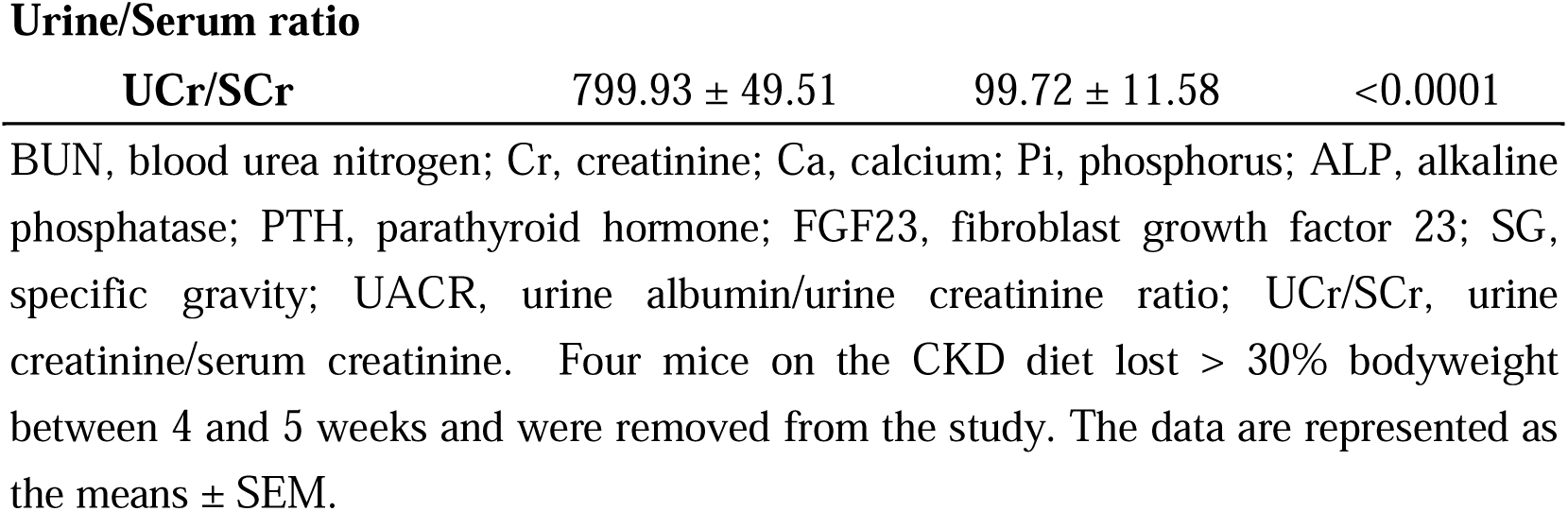
Body weight, serum, and urine biochemistries in control and CKD mice.

| Parameters | CONTROL | CKD | *P* Value |
| --- | --- | --- | --- |
|  | (n = 12) | (n = 8) |  |
| **Body Weight (g)** | 27.07 ± 1.81 | 19.06 ± 2.75 | <0.0001 |
| **Serum** |  |  |  |
| **BUN (mg/dL)** | 25.51 ± 1.18 | 65.16 ± 1.82 | <0.0001 |
| **Cr (mg/dL)** | 0.33 ± 0.00 | 0.58 ± 0.01 | <0.0001 |
| **Ca (mg/dL)** | 9.70 ± 0.15 | 10.65 ± 0.18 | <0.0001 |
| **Pi (mg/dL)** | 9.02 ± 0.24 | 13.32 ± 0.62 | <0.0001 |
| **ALP (IU/L)** | 162.92 ± 22.06 | 120.80 ± 6.01 | NS |
| **PTH (pg/mL)** | 1070.35 ± 154.40 | 1967.13 ± 204.40 | <0.01 |
| **FGF23 (ng/mL)** | 0.32 ± 0.05 | 47.74 ± 4.56 | <0.0001 |
| **Total P1NP**<br>**(ng/mL)** | 0.42 ± 0.08 | 0.78 ± 0.09 | <0.01 |
| **αCTx (ng/mL)** | 6.19 ± 1.86 | 16.45 ± 1.97 | <0.01 |
| **Urine** |  |  |  |
| **Cr (mg/dl)** | 262.50 ± 18.30 | 62.50 ± 8.18 | <0.0001 |
| **Albumin** | 141.25 ± 8.75 | 42.50 ± 8.18 | <0.0001 |
| **SG** | >1.04 ± 0.00 | 1.02 ± 0.00 | <0.0001 |
| Urine/Serum ratio |  |  |  |
| UCr/SCr | 799.93 ± 49.51 | 99.72 ± 11.58 | <0.0001 |

BUN, blood urea nitrogen; Cr, creatinine; Ca, calcium; Pi, phosphorus; ALP, alkaline phosphatase; PTH, parathyroid hormone; FGF23, fibroblast growth factor 23; SG, specific gravity; UACR, urine albumin/urine creatinine ratio; UCr/SCr, urine creatinine/serum creatinine. Four mice on the CKD diet lost > 30% bodyweight between 4 and 5 weeks and were removed from the study. The data are represented as the means ± SEM.

### 3.2. PHOSPHO1 and TNAP expression are altered in bones of CKD mice

*Phospho1* expression was increased and *Alpl* expression decreased in the femur of CKD mice when compared to control mice. The expression of *Enpp1, Slc20a2, Ank, Bglap, Pdpn, Runx2, Bmp2, and Tnfrsf11b* was decreased whereas femoral expression of *Fgf23*, and *Adipoq, were* increased in CKD mice when compared to control mice (Fig. 1A). The changes in *Phospho1* and *Alpl* expression in femurs of CKD mice were confirmed at the protein level (Figs. 1B, C).

**Fig 1.**
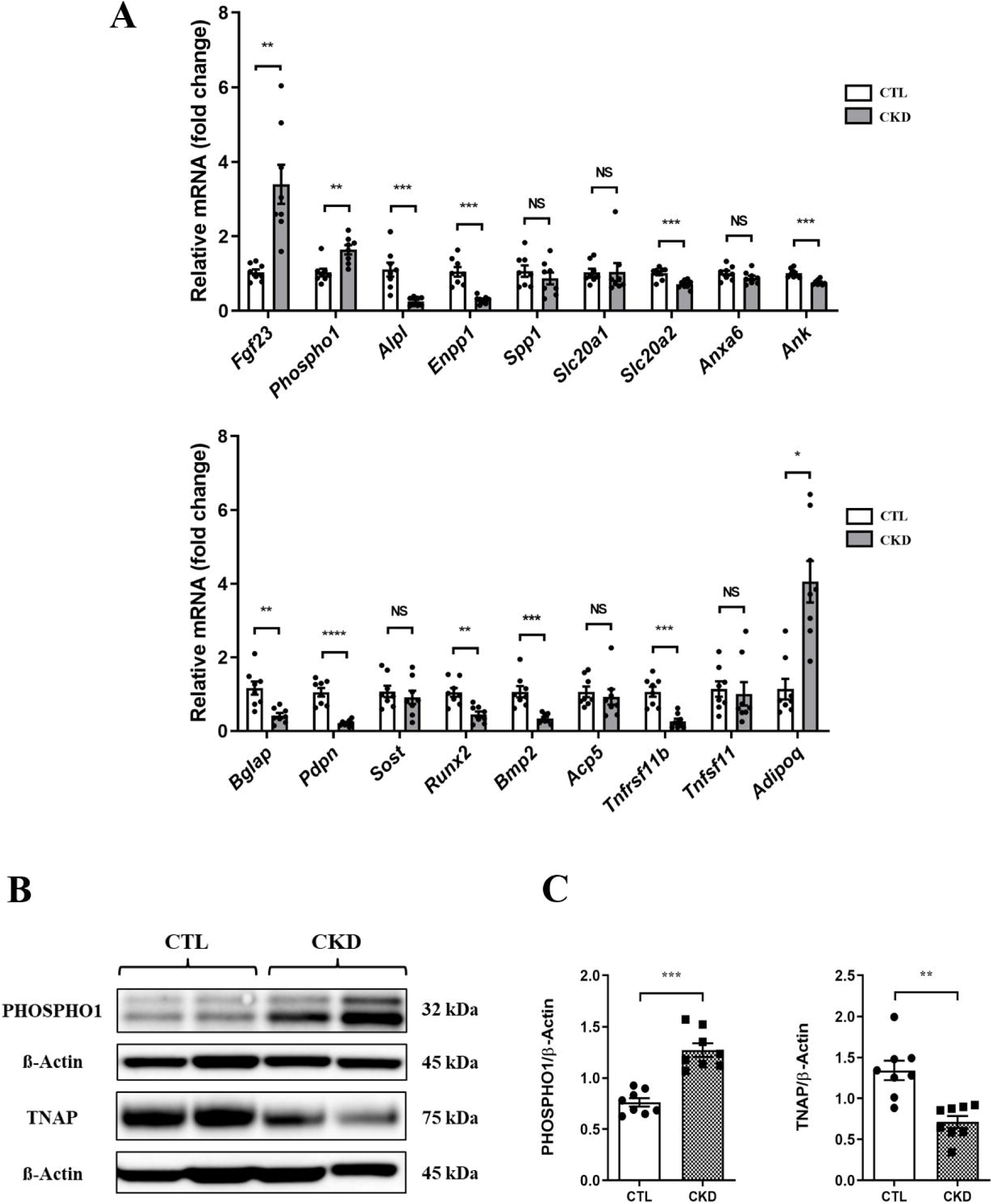
Expression of osteoblast and mineralization markers in mouse femurs from CTL and CKD mice. (A) *E*xpression of key mineralization and osteoblast marker genes in femurs of CTL and CKD mice at end of the study (13 weeks of age). Of note, *Fgf23* and *Phospho1* expression were increased and *Alpl* expression was decreased in the femurs of the CKD-MBD mice. (B) Representative image of 2 CTL and 2 CKD-MBD femurs analyzed by western blot for PHOSPHO1 and TNAP expression (C) Quantification of PHOSPHO1 and TNAP expression indicated that PHOSPHO1 was increased and TNAP was decreased in the femur of CKD-MBD mice compared with control mice. The data are represented as the mean ± SEM (n = 8); * *p* < 0.05; ** *p* < 0.01; *** *p* < 0.001; **** *p* < 0.0001).

### 3.3. Cortical BMD is increased in CKD mice and influenced by PHOSPHO1 status

Trabecular BMD, bone volume/tissue volume (BV/TV), thickness (Th), number (N), structural model index (SMI) and connectivity density (Conn Dn.) of the tibiae were all decreased in CKD mice when compared to controls (Fig. 2). Cortical bone parameters were also altered in CKD mice; cortical BMD was generally increased whereas cross-sectional area (CSA), cortical thickness, resistance to torsion, and Imin and Imax were all generally lower over the entire tibial length of CKD mice (Figs. 3a, c, d, f, i, j). Consistent with the thinner cortex the medullary area, the endosteal perimeter was increased and the periosteal perimeter decreased in the CKD mice (Figs. 3b, g, h). The histomorphometric analysis confirmed the reduced trabecular BV/TV in the CKD mice (Fig. S4A, i & ii, and Fig. S4B). The osteoid volume/bone volume (OV/BV) was increased in CKD mice confirming the impaired mineralization in this compartment (Fig. S4A, iii & iv, and Fig S4B). Osteoclast number associated with trabecular bone within the primary spongiosa of CKD mice was increased (Fig. S4A, v & vi and Fig. S4B); an observation consistent with decreased *Tnfrsf11b* (osteoprotegerin) expression in CKD bones (Fig. 1A) and higher serum αCTX concentrations in CKD mice (Table 1).

**Fig 2.**
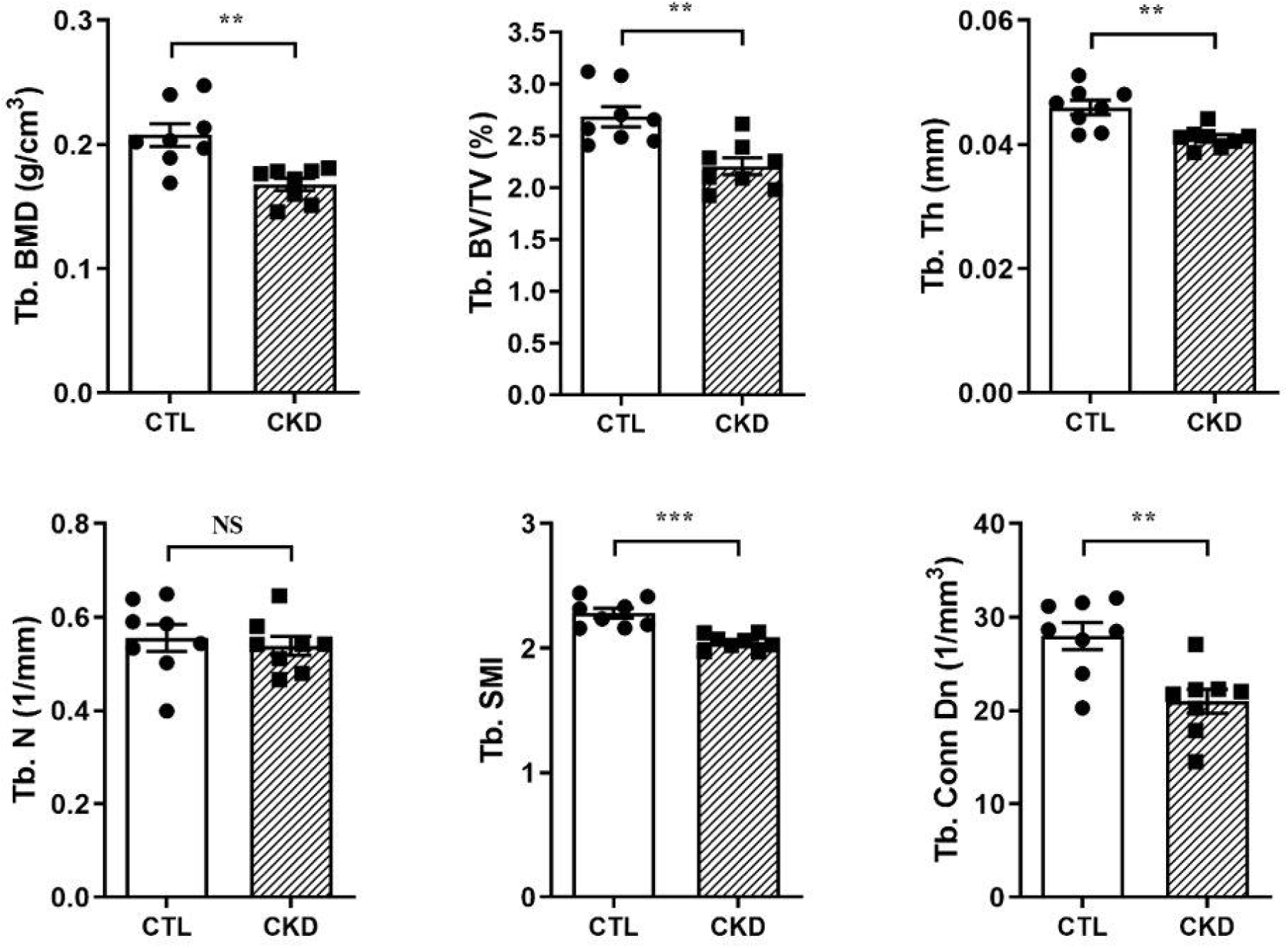
Micro-CT analysis of trabecular bone of the tibia. Micro-CT analysis of tibia from male C57BL/6 mice fed a CTL or CKD diet for 5 weeks. Tb. BMD (trabecular bone mineral density; g/cm^3^); Tb. BV/TV (trabecular bone volume/tissue volume; %); Tb. Th. (trabecular thickness; mm); SMI (structure model index); Tb. Conn Dn (trabecular connectivity density; mm^-3^) were all decreased in the CKD-MBD mice. Tb. N. (trabecular number; mm^-1^) was unchanged. Tibia of n = 8 (CTL mice) vs n = 8 (CKD-MBD mice) biological replicates were analysed. The data are represented as the means ± SEM. \**p* < 0.05; \*\**p* < 0.01; \*\*\**p* < 0.001 versus CTL.

**Fig 3.**
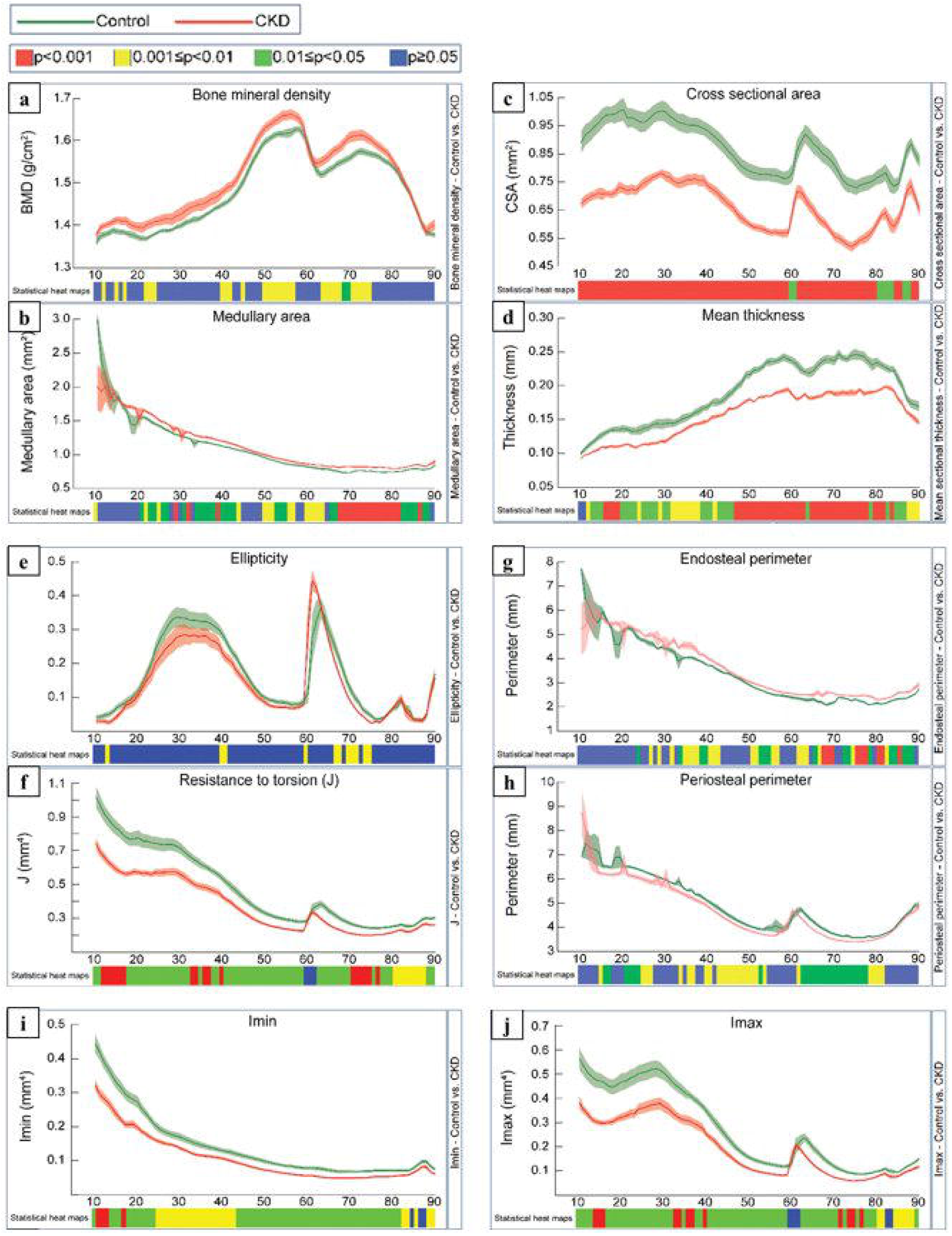
Micro-CT analysis of whole cortical bone of the tibia. Micro-CT analysis of tibia from male C57BL/6 mice fed a CTL or CKD diet for 5 weeks. Quantification of whole bone analyses of cortical bone between 10% and 90% of total tibial length, excluding proximal and distal metaphyseal bone, of CTL and CKD tibia at 13 weeks of age. (a) BMD (bone mineral density; g/cm^3^), (b) medullary area (cm^2^) and (g) endosteal perimeter (mm) were generally increased and (c) CSA (cross-sectional area; mm^2^), (d) mean thickness (mm), (f) resistance to torsion (J; mm^4^), (h) periosteal perimeter (mm), (i) Imin (mm^4^), (j) Imax (mm^4^) were generally decreased in the CKD-MBD bones. Tibia of n = 8 (CTL mice) vs n = 8 (CKD mice) biological replicates were analysed. *p* < 0.05 was significant and *p* ≤ 0.01–0.05 was noted as green, *p* ≤ 0.001–0.01 as yellow and *p* ≤ 0.000–0.001 as red. Not significant is noted as blue.

The increased cortical BMD in CKD-MBD mice (Fig. 3a) aligns with the higher PHOSPHO1 expression in the cortical bone shafts, despite being an unexpected finding in the CKD-MBD mice (Figs. 1A-C). To explore this further, we next examined bone from PHOSPHO1 deficient (P1KO) mice maintained on the 0.2% adenine supplemented diet for 5 weeks. Cortical analysis was performed on datasets derived from μCT scan images at 50% of the total tibial length as this region of bone from CKD mice had a higher BMD than control counterparts (Fig 3a). As previously noted, (Fig. 3a), the cortical BMD of WT CKD mice was increased compared to WT control mice but, in contrast, no such increase was apparent in P1KO CKD mice, which had a BMD similar to their respective P1KO controls but as expected lower than the BMD of WT control mice (Fig. 4). Structural cortical bone changes were also influenced by the absence of PHOSPHO1 in the P1KO mice; the CKD-induced increases in porosity and decreases in BV/TV, CSA, and Th noted in WT CKD mice were all blunted in P1KO CKD mice compared to P1KO control mice (Fig. 4).

**Fig 4.**
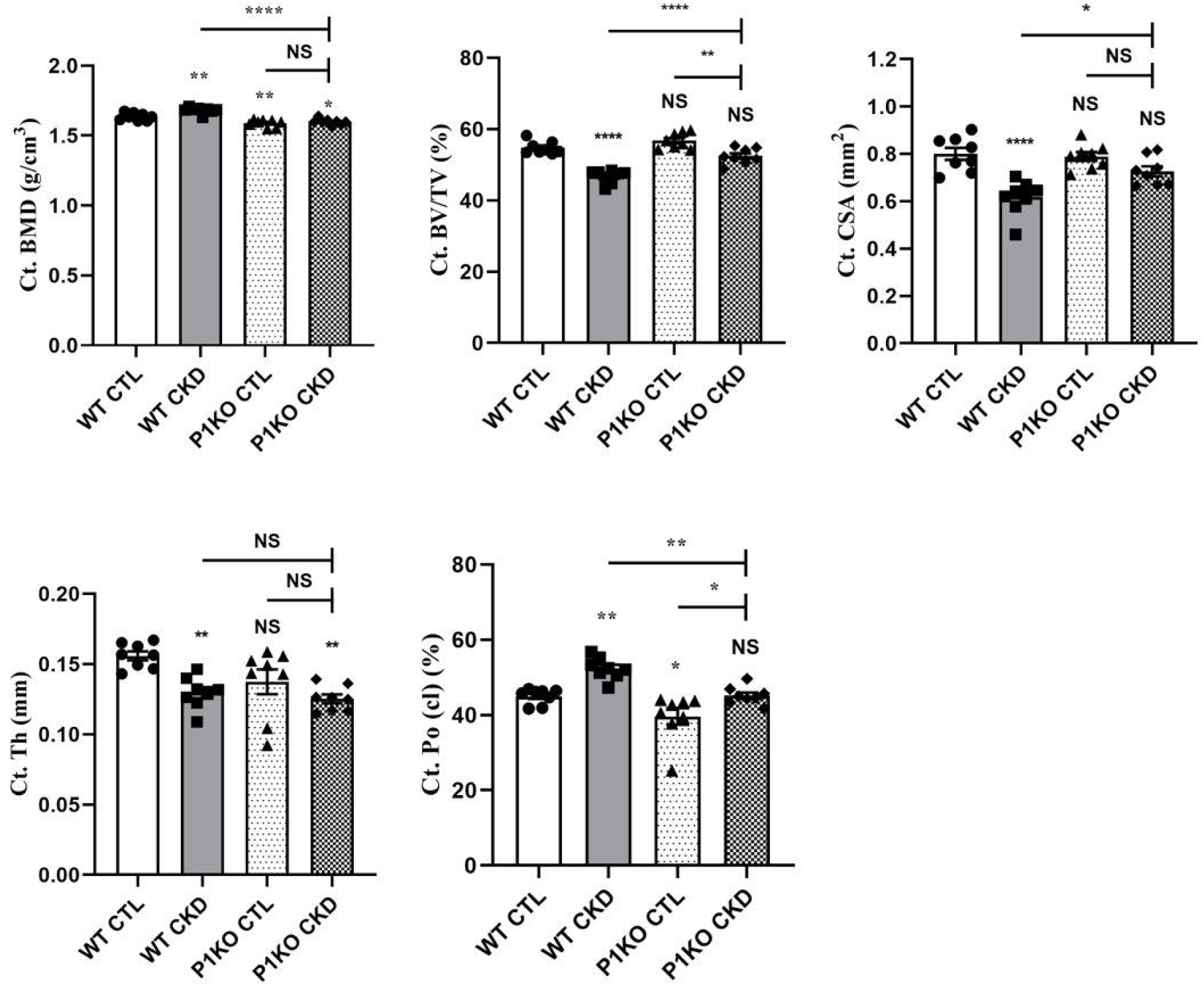
Micro-CT analysis of cortical bone of wild-type (WT) and PHOSPHO1 deficient CTL and CKD mice. Quantification of cortical bone mineral density (Ct. BMD), cortical bone volume/tissue volume (Ct. BV/TV), cortical cross-sectional area (Ct. CSA), cortical thickness (Ct. Th), and closed pore porosity (Ct Po (cl)) at 50% of the total tibial length from the top of the tibia. Of note, BMD was increased in the WT CKD-MBD tibia but not the PHOSPHO1 deficient CKD-MBD tibia when compared to their respective controls. The data are represented as the mean ± SEM (n = 8); * *p* < 0.05; ** *p* < 0.001; **** *p* < 0.0001.

### 3.4. Pi, PTH, and FGF23 perturb ECM mineralization and the expression of key mineralization markers in primary osteoblasts

To determine the causes of the mineralization defects noted in the CKD mice, we investigated the direct effects of FGF23, PTH, and Pi on the expression of PHOSPHO1 and TNAP and other key regulators of mineralization by primary osteoblasts in cultures. Over 28 days, the basal Pi-substrate free mineralization media promoted matrix mineralization (Figs. S5A, B) and PHOSPHO1 and TNAP expression in a temporal manner at both the gene and protein level confirming the suitability of this culture model for our purposes (Figs. S5C-E).

At concentrations of 2 mM and above, Pi significantly down-regulated *Phospho1, Alpl, and Bglap* mRNA expression (*p* < 0.01, Fig. 5A). In contrast, *Enpp1, Spp1* and *Slc20a1* expression was increased at the higher Pi concentrations (*p* < 0.05, Fig. 5A). Cell viability as assessed by Alamar blue and LDH release was unaffected at all Pi concentrations tested (Fig. S6). PHOSPHO1 and TNAP protein expression were also inhibited by increasing Pi concentrations whereas the addition of Pi, 3 mM and above, increased the formation of mineralized bone nodules in a dose-dependent manner (*p* < 0.001, Figs. 5B, C).

**Fig 5.**
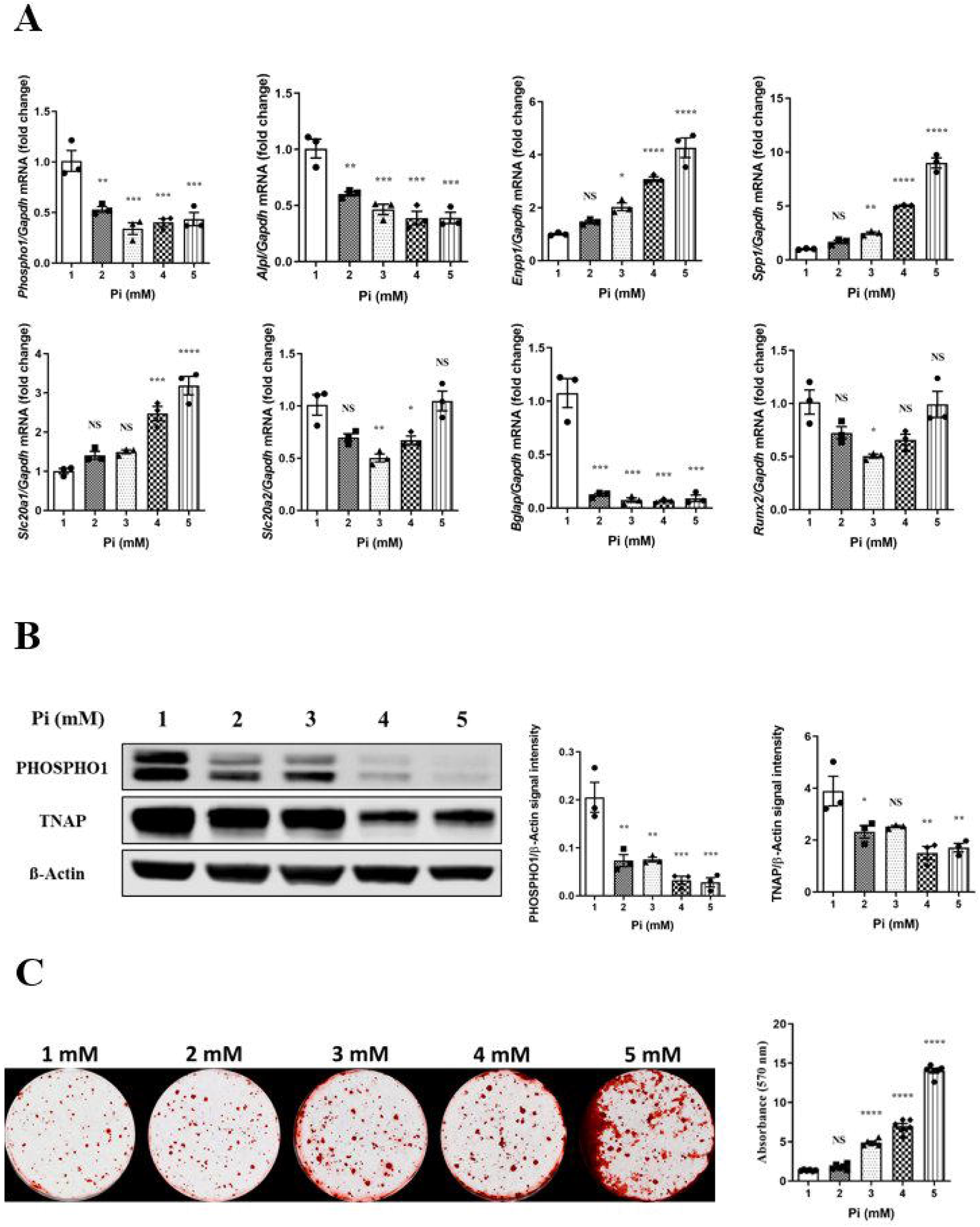
Regulation of key mineralization associated genes, proteins and osteoblast extracellular matrix mineralization by Pi in primary osteoblasts. (A) Expression analysis of *Phospho1, Alpl, Enpp1, Spp1, Slc20a1, Slc20a2, Bglap, and Runx2* by osteoblasts in response to Pi (1-5 mM), (B) western blotting analysis and quantification of PHOSPHO1 and TNAP expression in response to Pi and (C) representative images and quantification of alizarin red staining in response to Pi for 28 days after confluency. PHOSPHO1 and TNAP at the gene and protein level were decreased with increasing Pi concentrations whereas matrix mineralization increased with increasing Pi concentrations. The data are represented as the mean ± SEM (n = 3); \**p* < 0.05; ** *p* < 0.01; *** *p* < 0.001; **** *p* < 0.0001.

Administration of PTH at > 5 nM downregulated the expression *of Phospho1, Alpl* and *Bglap* (*p* < 0.01, Fig. 6A). *Enpp1, Slc20a2*, and *Runx2* expression were also decreased but only at higher PTH concentrations (*p* < 0.05, Fig. 6A). Reduction of PHOSPHO1 and TNAP protein expression by increasing PTH concentrations mirrored the changes in gene expression (Fig. 6B). The addition of PTH inhibited ECM mineralization and this was noted at concentrations as low as 0.5 nM. Mineralization was completely abolished at 25 and 50 nM (*p* < 0.001, Fig. 6C).

**Fig 6.**
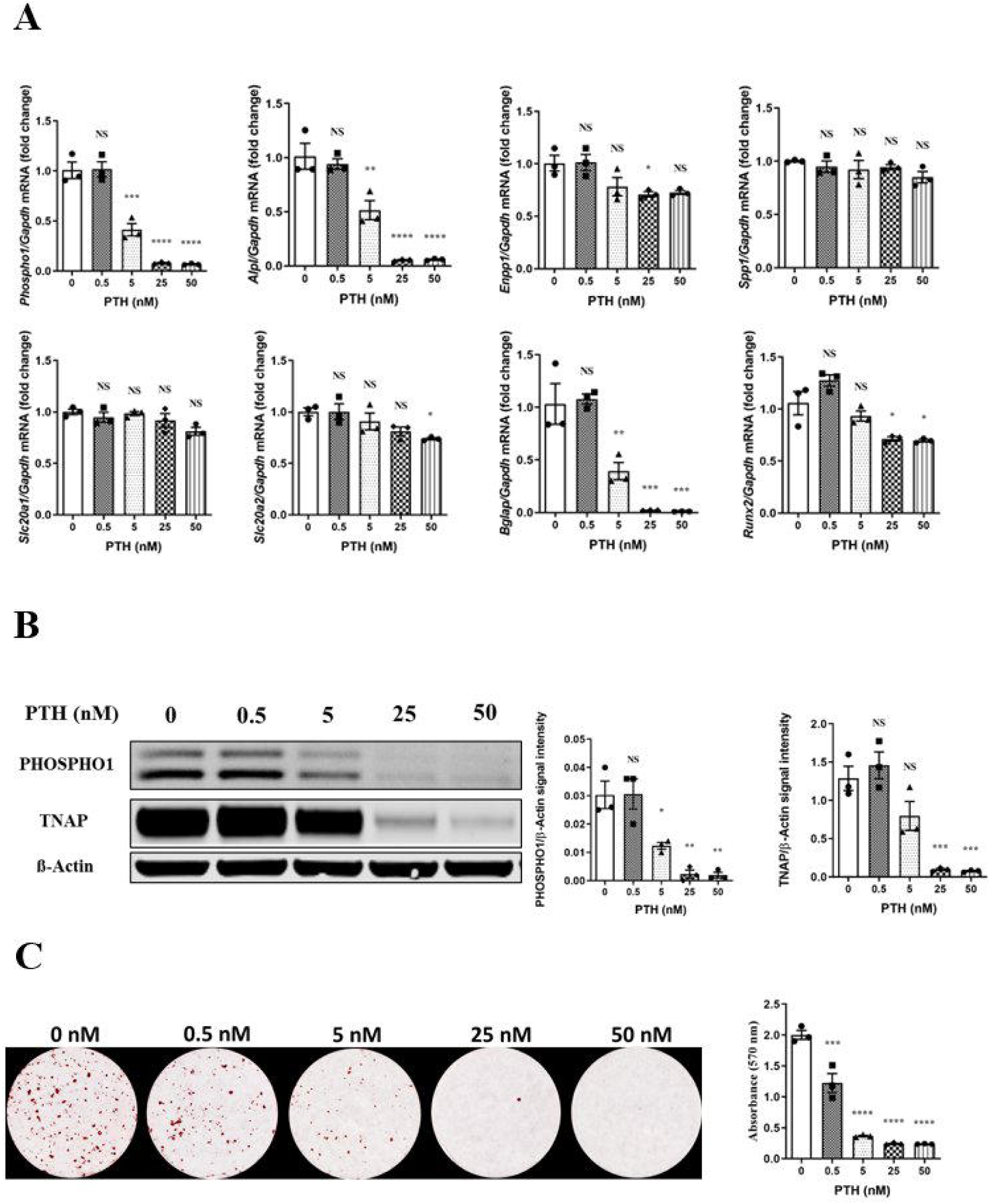
Regulation of key mineralization associated genes, proteins and osteoblast extracellular matrix mineralization by PTH in primary osteoblasts. (A) Expression analysis of *Phospho1, Alpl, Enpp1, Spp1, Slc20a1, Slc20a2, Bglap, and Runx2* by osteoblasts in response to PTH (0-50 nM), (B) western blotting analysis and quantification of PHOSPHO1 and TNAP expression in response to PTH and (C) representative images and quantification of alizarin red staining in response to PTH for 28 days after confluency. PHOSPHO1 and TNAP at the gene and protein level and matrix mineralization were all decreased with increasing Pi concentrations. The data are represented as the mean ± SEM (n = 3); \**p* < 0.05; ** *p* < 0.01; *** *p* < 0.001; **** *p* < 0.0001.

Exposure to FGF23 had little effect on the expression of the genes studied although both *Phospho1* and *Alpl* expression were decreased but only at the highest FGF23 concentrations (*p* < 0.05, Fig. 7A). The addition of Klotho to the FGF23 supplemented cultures had no further effects on gene expression when compared with FGF23 alone (data not shown). A similar trend was also noted at the protein level where PHOSPHO1 and TNAP expression decreased in a FGF23 concentration-dependent manner but this change did not reach statistical significance from control-treated cultures (Fig. 7B). A similar response was observed in the presence of FGF23 and Klotho (data not shown). FGF23 with or without klotho had no effects on ECM mineralization of primary osteoblasts at the concentrations tested (Fig. 7C and data not shown).

**Fig 7.**
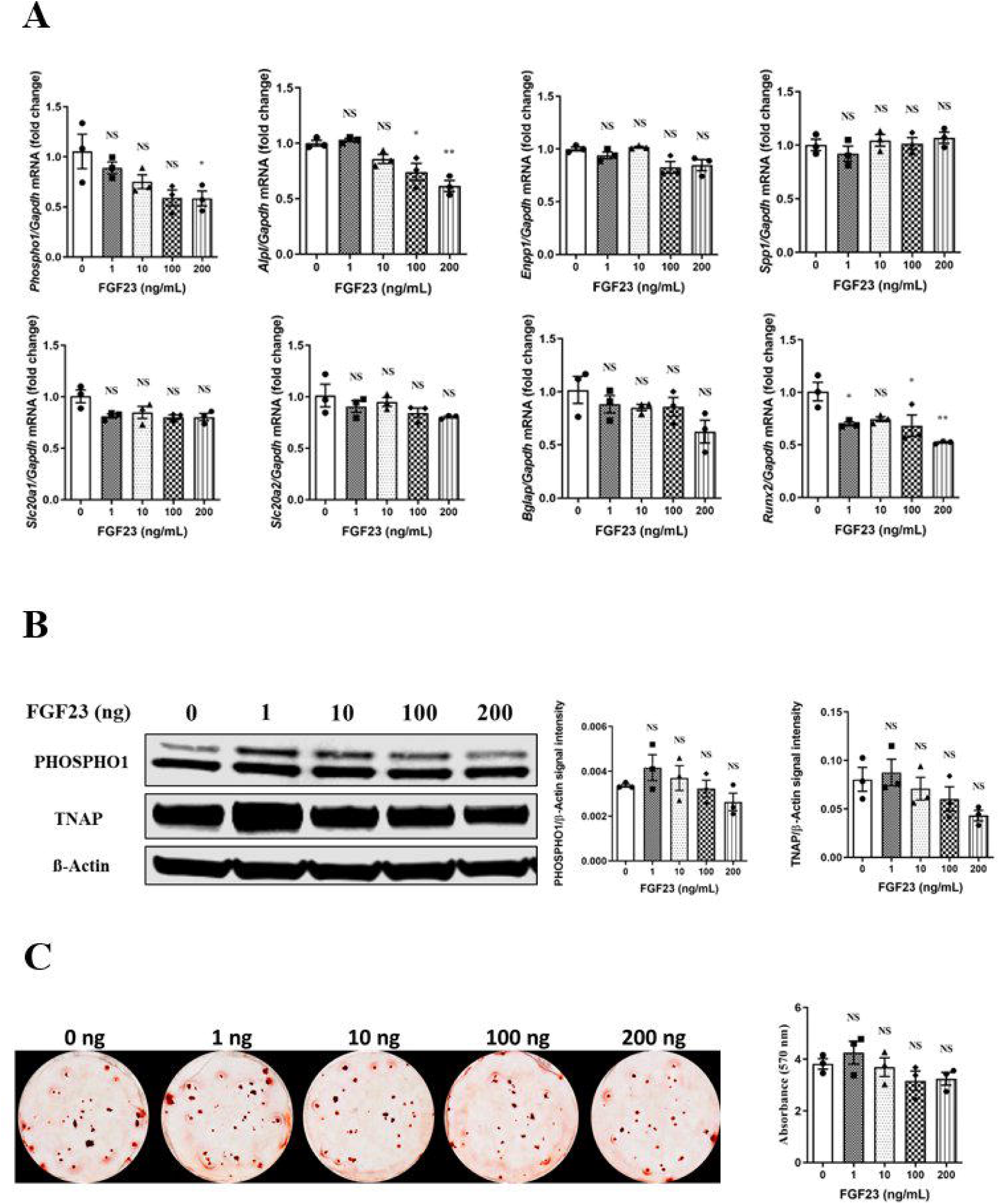
Regulation of key mineralization associated genes, proteins and osteoblast extracellular matrix mineralization by FGF23 in primary osteoblasts. (A) Expression analysis of *Phospho1, Alpl, Enpp1, Spp1, Slc20a1, Slc20a2, Bglap, and Runx2* by osteoblasts in response to FGF23 (0-200 ng/ml), (B) western blotting analysis and quantification of PHOSPHO1 and TNAP expression in response to FGF23 and (C) representative images and quantification of alizarin red staining in response to FGF23 for 28 days after confluency. *Phospho1* and *Alpl* gene expression were decreased at the highest FGF23 concentrations but non-significant differences were noted with PHOSPHO1 and TNAP expression and matrix mineralisation. The data are represented as the mean ± SEM (n = 3); \**p* < 0.05; ** *p* < 0.01; *** *p* < 0.001; **** *p* < 0.0001.

## 4. Discussion

This study has shown that PHOSPHO1 and TNAP, two phosphatases required for bone mineralization, have altered expression in ROD. Specifically, the ROD phenotype was characterized by increased cortical BMD and this response may be mediated by increased PHOSPHO1 expression. However, the altered PHOSPHO1 expression is unlikely to be a direct result of the altered endocrine milieu as PTH, FGF23 and Pi all decreased PHOSPHO1 expression in osteoblast cultures. Overall, utilizing *in vivo* and *in vitro* experimental models of CKD, this study is the first to implicate PHOSPHO1 function in the altered mineralization status of bones in an animal model of ROD.

In humans, deteriorating renal function contributes to the progression of ROD which results in bone loss, osteoporosis, and eventually increased morbidity and mortality resulting from fractures and/or cardiovascular disease (Gal-Moscovici and Sprague 2007). A similar bone phenotype was mirrored in this present study where cortical thinning, lower BV/TV, and increased cortical porosity were noted in the adenine-fed mice. The loss of bone is likely to be multi-factorial but PTH enhanced bone resorption via altered RANKL and OPG expression is likely to predominate (Ma et al. 2001). In the early stages of CKD, the low bone turnover disease results from bone cell inactivity due to PTH resistance, as well as reduced calcitriol levels, and accumulation of uremic toxins (Couttenye et al. 1999). When renal function further deteriorates, the chronically increased PTH levels overcome peripheral PTH resistance and activate the indolent bone cells, leading to high turnover bone disease (Drüeke and Massy 2016). Bone resorption predominates in both high and low bone turnover disease and the resultant elevated serum Ca and Pi levels promote bone extra-skeletal (vascular) calcification (Zheng et al. 2016). In agreement with the results of this present study, others have also reported increased cortical porosity and compromised bone architecture in CKD animal models although inconsistent effects on the cortical and trabecular compartments have been reported (Jia et al. 2013; Metzger et al. 2021; Miller et al. 1998; Ogirima et al. 2006). Although patients with CKD have been reported to have lower cortical BMD inconsistencies in trabecular and cortical BMD in CKD animal models also exist (Lau et al. 2013b; Nickolas et al. 2013). Specifically, in a mouse nephrectomy model in which serum Pi levels were unchanged, trabecular and cortical BMD were increased and decreased, respectively which was the opposite to that found in this present study (Lau et al. 2013b). The increased trabecular BMD was not influenced by dietary phosphate content whereas the decreased cortical BMD was only noted in mice fed a high phosphate (0.9%) and not a normal phosphate (0.5%) containing diet (Lau et al. 2013b). In this present study, mice were fed 0.9% phosphate-containing diet and analysis revealed that at no location along the entire cortical bone shaft was BMD lower in the CKD mice. The spectrum of bone phenotypes reported in CKD-MBD models may reflect the differing serum PTH levels at the point of the study, as progressive SHPT is linked with different effects on bone quality and structure (Miller et al. 1998). Furthermore, whether differential expression of PHOSPHO1 and TNAP within the trabecular and cortical bone compartments contributes to the divergent BMD response is unclear and requires further investigation.

The high bone turnover status in SHPT will contribute to bone that is less mineralized, a hallmark of stage 4 and 5 CKD, and lead to reduced mechanical strength and increased risk of fractures (Drüeke and Massy 2016). Similarly, in this present study, PTH-induced skeletal remodeling is likely to, at least in part, explain the poorly mineralized trabecular bone noted in this study although PTH exposure can also inhibit osteoblast differentiation and thus indirectly delay osteoid production and matrix mineralization (Qin et al. 2004). Additionally, the CKD-driven increase in osteocyte secretion of Wnt/β-catenin□signaling inhibitors, such as FGF23, dickkopf 1, and sclerostin may negatively affect osteoblast function and contribute to the mineralization defect in ROD (Evenepoel et al. 2015; Murali et al. 2016a). The results of this present study offer changes to PHOSPHO1 and TNAP osteoblast expression as an additional/alternative explanation for the altered bone mineralization status associated with ROD.

Monitoring serum ALP has been regarded as a useful serum marker of bone turnover in ROD however its expression in bone, functioning as a phosphatase capable of mineralizing osteoid has not to our knowledge been explored in the pathogenesis of ROD (Bervoets et al. 2003). The decreased *Alpl* expression in CKD cortical bone was not however consistent with the observed increased cortical BMD and we hypothesize that the latter is possibly driven by increased PHOSPHO1 expression (Huesa et al. 2015). To examine this further we determined cortical BMD and other structural parameters in P1KO CKD mice. In the absence of PHOSPHO1, cortical BMD in control mice was decreased as previously reported and the increased BMD in cortical bone of CKD wild-type mice was not observed in the P1KO CKD mice (Yadav et al. 2011). Furthermore, other structural parameters such as cortical porosity, thickness and CSA were also improved in P1KO CKD mice. Whilst supportive of our hypothesis, the mechanisms responsible for the increased PHOSPHO1 in cortical bone are unclear and cannot be explained by the direct effects of Pi, FGF23, and PTH which are all inhibitory to PHOSPHO1 expression by osteoblasts *in vitro* as shown in this study. A compensatory mechanism in an attempt to protect the bone from hypomineralization may be a possibility but further work on this and whether PHOSPHO1 deficiency improves bone health in ROD by decreasing cortical porosity is warranted (Metzger et al. 2020). The decreased cortical bone TNAP expression could be a direct effect of Pi and PTH on osteoblasts as shown by the *in vitro* data of this and other studies (Houston et al. 2016; Rendenbach et al. 2014). Furthermore, whilst not observed in this study, others have reported a direct inhibitory effect of FGF23 on osteoblast matrix mineralization which may be mediated by decreased TNAP expression and an accumulation of PPi, a potent inhibitor of mineralization (Murali et al. 2016a; Shalhoub et al. 2011). However, indirect systemic effects via disrupted vitamin D status and Pi and Ca metabolism are also likely to contribute to the altered TNAP expression in bone of CKD-MBD mice (Bover et al. 2018; Rendenbach et al. 2014).

Several studies have reported that mineralizing cells including odontoblasts, chondrocytes, and osteoblasts are sensitive to P_i_ and respond by altering the expression of mineralization-associated genes and transcription factors (Foster et al. 2006; Khoshniat et al. 2011). The regulation of biomineralization by Pi may be related to its ability to stimulate MV release and/or the accumulation of type III NaP(i) transporter (PiT1) in osteoblasts promoting Pi uptake and ECM mineralization (Foster et al. 2006; Yoshiko et al. 2007). In relation to this present study, the availability of exogenous Pi to promote osteoblast matrix mineralization by-passes the requirement for Pi production from phosphocholine and phosphoethanolamine by PHOSPHO1 and PPi by TNAP (Ciancaglini et al. 2010; Houston et al. 2004; Roberts et al. 2007) and may explain the concentration-dependent decrease in PHOSPHO1 and TNAP expression by exogenous Pi as part of a negative feedback mechanism (Foster et al. 2006). In this regard, human PHOSPHO1 shares approximately 30% homology at the amino acid level with a tomato phosphate starvation-induced gene product, LePS2, which possesses phosphatase activity that can convert organic phosphorus into available Pi. Intriguingly, LePS2 expression is tightly and negatively regulated by Pi availability and is thus induced in the absence, but repressed in the presence of Pi (Stenzel et al. 2003). It is unknown if such a Pi negative feedback mechanism controls PHOSPHO1 expression but the increased osteoclast resorption observed in ROD will bring about the release of Pi which will contribute to the observed hyperphosphatemia and impede the skeleton from exerting its normal reservoir function when serum Pi concentrations increase (Hruska et al. 2008). In such a scenario, the resulting Pi stress conditions experienced by the skeleton may drive higher PHOSPHO1 expression in a similar way to the LePS2 protein and other phosphatases such as OsACP1 a PHOSPHO1-like acid phosphatase in rice (Deng et al. 2022).

In summary, this study has identified PHOSPHO1 as a possible mediator in the development of the cortical bone phenotype in ROD, thus providing a foundation for future research to explore potential therapies to improve bone health in CKD-MBD.

## CRediT authorship contribution statement

**Shun-Neng Hsu:** Conceptualization, Formal Analysis, Methodology, Investigation, Writing – Original draft, Funding acquisition. **Louise A Stephen:** Formal Analysis, Methodology, Investigation, Supervision, Writing – Review & Editing. **Scott Dillon:** Formal Analysis, Methodology, Investigation. **Elspeth Milne:** Formal Analysis, Methodology. **Behzad Javaheri:** Formal Analysis, Methodology. **Andrew A Pitsillides:** Methodology, Investigation. **Amanda Novak:** Conceptualization, Methodology. **Jose Luis Millán:** Investigation. **Vicky E Macrae:** Conceptualization, Supervision, Funding acquisition; Writing – Review & Editing. **Katherine A Staines:** Conceptualization, Investigation, Supervision, Writing – Review & Editing, Funding acquisition. **Colin Farquharson:** Conceptualization, Investigation, Writing – Review & Editing, Supervision, Funding acquisition. All authors approved the final version of the manuscript.

## Transparency document

The Transparency document associated with this article can be found, in online version.

## Declaration of competing interest

The authors declare that they have no competing interest.

### Ethical Approval

All experimental protocols were approved by Roslin Institute’s Animal Users Committee and the animals were maintained in accordance with UK Home Office guidelines for the care and use of laboratory animals, and with the ARRIVE guidelines.

## Acknowledgments

We thank Elaine Seawright and Yao-Tang Lin for technical assistance; Darren Smith, Tricia Mathieson, Heather Warnock, Lorraine Blackford, Lee Mcmanus, and the staff of the animal care facility at Biological Research Facility (BRF) providing animal support; Colin Wood and the staff at Easter Bush Pathology, Royal (Dick) School of Veterinary Studies, University of Edinburgh for conducting mouse serum biochemistries.

## Supplementary figure legends

**Fig S1. Schematic view of the 5-week adenine induced CKD-MBD model and time-dependent changes in body weight**. (A) Eight-week-old C57BL/6 male mice were randomly allocated to either a control (CTL; n=12) or CKD (n = 12) group. Mice in the CKD-MBD group were fed a casein-based diet containing 0.2% adenine diet for 5 weeks. Mice in the CTL group were fed a casein-based diet without adenine. (B) The bodyweight of the CKD-MBD mice progressively decreased during the 5-weeks on the adenine supplemented diet. The data are represented as the means ± SEM. **** *p* < 0.0001 as compared to CTL mice of the same age. Note, 4 mice in the CKD-MBD group lost more than 30% of their body weight and were removed from the study.

**Fig S2. Characterization of renal pathology in CKD mouse model**. (A) Representative photomicrographs of hematoxylin and eosin (H&E; i-iv), Masson’s trichrome (MT; v & vi), and von Kossa (VK; vii & viii) stained kidney sections from CTL and CKD mice at end of the study (13 weeks of age). (i & ii) kidney sections showing gross pathology, (d) atrophic tubuli with protein casts (green arrows) and dilated Bowman’s space (blue arrow). (f) Dilated tubules (green arrows) and interstitial fibrosis (blue arrows. (h) Calcification of tubular structures (blue arrows). Scale bar, (i & ii) 500 μm, (iii-viii) 100 μm. (B) Renal scoring of tubular atrophy, protein casts, interstitial inflammation, and renal fibrosis of sections from 4 CTL and 4 CKD mice. All indices were higher in kidneys from CKD-MBD mice. Renal scoring scale: 0 = normal; 1 = mild, involvement of <25% of the cortex; 2 = moderate, involvement of 25 to 50% of the cortex; 3 = severe, involvement of 50 to 75% of the cortex; 4 = extensive damage involving >75% of the cortex. The data are represented as the mean ± SEM (n = 4); **** *p* < 0.0001.

**Fig S3. Expression levels of injury associated markers in kidneys of CTL and CKD mice. (A)** *Fgf23, Spp1 and Lcn2* expression was higher whereas *Slc34a1* was lower in in kidneys of CKD-MBD mice at end of the study (13 weeks of age). Four random samples from each of the CTL and CKD groups were selected for analysis. (B) Representative western blot of osteopontin (OPN) and type II sodium-phosphate cotransporter (NaPi-2a) protein expression from BBMV of kidneys at end of study (13 weeks of age). The increased osteopontin and decreased NaPi-2a in CKD-MBD kidneys confirm gene expression data. The data are represented as the mean ± SEM (n = 4); * *p* < 0.05; ** *p* < 0.01; *** *p* < 0.001; **** *p* < 0.0001

**Fig S4. Histological characterization of trabecular bone in CKD mice (A)** Representative photomicrographs of tibia sections stained for hematoxylin and eosin (H&E; i & ii) and Goldner’s trichrome (iii & iv) and reacted for tartrate acid phosphatase activity (TRAP; v & vi; blue arrow) from CTL and CKD-MBD mice at end of study (13 weeks of age). Scale bar, 100 μm. (B) Bone volume/tissue volume (BV/TV) was decreased in CKB-MBD mice whereas osteoid volume/bone volume (OV/BV); osteoclast surface/bone surface (Oc.S/BS); number of osteoclasts/bone surface (N.Oc/BS) were all increased in CKD-MBD mice. The data are represented as the mean ± SEM (n = 6); * *p* < 0.05; ** *p* < 0.01.

**Fig S5. Characterization of osteoblast culture model showing temporal increases in extracellular matrix mineralization and PHOSPHO1 and TNAP expression**. (A) Alizarin red staining, (B) quantification of matrix mineralization (C) RT-qPCR analysis of *Phospho1*, and *Alpl* mRNA expression, (D) western blot analysis, and (E) quantification of PHOSPHO1, and TNAP expression and by primary osteoblasts cultured in the basal Pi-substrate free mineralization medium over a 28-day culture period. The data are represented as the mean ± SEM (n = 3); * *p* < 0.05; ** *p* < 0.01; *** *p* < 0.001; **** *p* < 0.0001 in comparison with day 0, post confluence cultures.

**Fig S6. The effect of Pi on osteoblast viability**. Cells were exposed to Pi (1-5 mM) for 28 days after confluency and viability were assessed by (A) Alamar Blue assay, and (B) LDH assay. Cell viability was not affected by Pi at all concentrations tested. The data are represented as the mean ± SEM (n ≥ 3); NS, not significance from the 1 mM control group.

## Supplementary tables

**Table S1.**
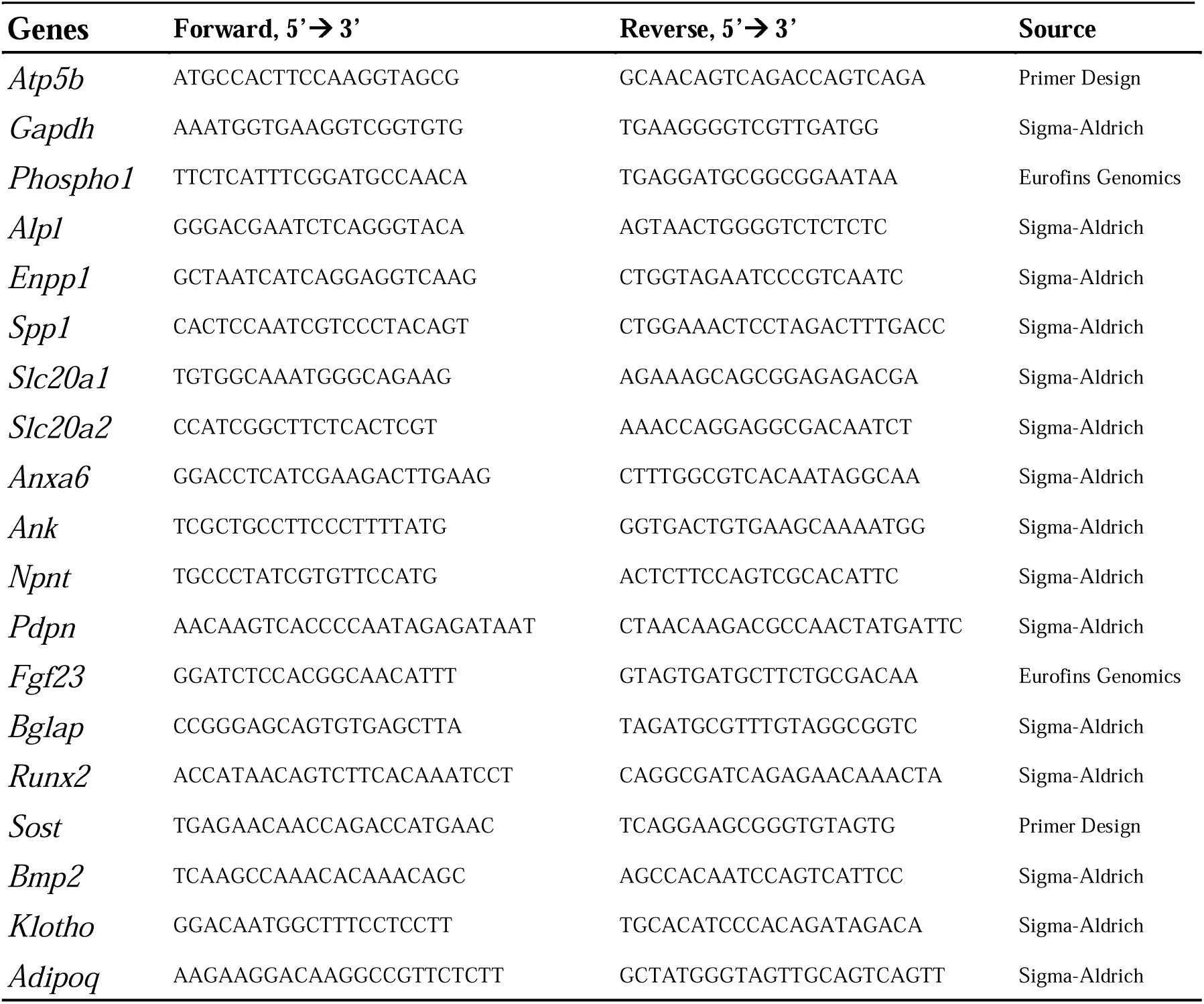
Sequences of primers used for qPCR.

**Table S2.** Primary antibodies used for western blotting.

| **Target** | **Source** | **Company** | **Dilution** | **Molecular weight** |
| --- | --- | --- | --- | --- |
| β-actin | Rabbit | Cell Signaling Technology | 1:4000 | 45 kDa |
| PHOSPHO1 | Human | AbD Serotec and Bio-Rad | 1:500 | 32 kDa |
| TNAP | Rat | R&D Systems | 1:1000 | 75 kDa |

**Table S3.** Secondary antibodies used for western blotting.

| **Target** | **Source** | **Company** | **Dilution** |
| --- | --- | --- | --- |
| β-actin | Goat anti-rabbit | Dako | 1:1000 |
| PHOSPHO1 | Goat anti-human | Bio-Rad | 1:1000 |
| TNAP | Goat anti-rat | R&D Systems | 1:1000 |
| β-actin | Donkey anti-rabbit | LI-COR Biosciences | 1:15000 |
| PHOSPHO1 | Goat anti-human | LI-COR Biosciences | 1:15000 |
| TNAP | Goat anti-rat | LI-COR Biosciences | 1:15000 |

## References

Bajwa, NM, Sanchez, CP, Lindsey, RC, et al. (2018) Cortical and trabecular bone are equally affected in rats with renal failure and secondary hyperparathyroidism. BMC Nephrol 19:24. DOI: 10.1186/s12882-018-0822-8.

Bervoets, ARJ, Spasovski, GB, Behets, GJ, et al. (2003) Useful biochemical markers for diagnosing renal osteodystrophy in predialysis end-stage renal failure patients. Am J Kidney Dis 41:997–1007. DOI: 10.1016/s0272-6386(03)00197-5.

Bover, J, Ureña, P, Aguilar, A, et al. (2018) Alkaline Phosphatases in the Complex Chronic Kidney Disease-Mineral and Bone Disorders. Calcif Tissue Int 103:111–124. DOI: 10.1007/s00223-018-0399-z.

Boyde, A, Staines, KA, Javaheri, B, et al. (2017) A distinctive patchy osteomalacia characterises Phospho1-deficient mice. Journal of anatomy 231:298–308. DOI: 10.1111/joa.12628.

Chande, S & Bergwitz, C (2018) Role of phosphate sensing in bone and mineral metabolism. Nature Reviews Endocrinology 14:637–655. DOI: 10.1038/s41574-018-0076-3.

Ciancaglini, P, Yadav, MC, Simão, AM, et al. (2010) Kinetic analysis of substrate utilization by native and TNAP-, NPP1-, or PHOSPHO1-deficient matrix vesicles. J Bone Miner Res 25:716–23. DOI: 10.1359/jbmr.091023.

Couttenye, MM, D ‘Haese, PC, Verschoren, WJ, et al. (1999) Low bone turnover in patients with renal failure. Kidney Int 56:S70–S76. DOI: https://doi.org/10.1046/j.1523-1755.1999.07308.x.

Deng, S, Li, J, Du, Z, et al. (2022) Rice ACID PHOSPHATASE 1 regulates Pi stress adaptation by maintaining intracellular Pi homeostasis. Plant, Cell & Environment 45:191–205. DOI: https://doi.org/10.1111/pce.14191.

Dillon, S, Staines, KA, Millán, JL, et al. (2019) How To Build a Bone: PHOSPHO1, Biomineralization, and Beyond. JBMR Plus 3:e10202. DOI: 10.1002/jbm4.10202.

Drüeke, T. & Massy, ZA (2016) Changing bone patterns with progression of chronic kidney disease. Kidney Int 89:289–302. DOI: 10.1016/j.kint.2015.12.004.

Evenepoel, P, D ‘Haese, P & Brandenburg, V (2015) Sclerostin and DKK1: new players in renal bone and vascular disease. Kidney Int 88:235–40. DOI: 10.1038/ki.2015.156.

Fang, Y, Ginsberg, C, Sugatani, T, et al. (2014) Early chronic kidney disease-mineral bone disorder stimulates vascular calcification. Kidney Int 85:142–50. DOI: 10.1038/ki.2013.271.

Foster, BL, Nociti, FH, Jr., Swanson, EC, et al. (2006) Regulation of cementoblast gene expression by inorganic phosphate in vitro. Calcif Tissue Int 78:103–12. DOI: 10.1007/s00223-005-0184-7.

Gal-Moscovici, A & Sprague, SM (2007) Bone health in chronic kidney disease-mineral and bone disease. Adv Chronic Kidney Dis 14:27–36. DOI: 10.1053/j.ackd.2006.10.010.

Graciolli, FG, Neves, KR, Barreto, F, et al. (2017) The complexity of chronic kidney disease-mineral and bone disorder across stages of chronic kidney disease. Kidney Int 91:1436–1446. DOI: 10.1016/j.kint.2016.12.029.

Hessle, L, Johnson, KA, Anderson, HC, et al. (2002) Tissue-nonspecific alkaline phosphatase and plasma cell membrane glycoprotein-1 are central antagonistic regulators of bone mineralization. Proc Natl Acad Sci U S A 99:9445–9. DOI: 10.1073/pnas.142063399.

Houston, B, Stewart, A. & Farquharson, C (2004) PHOSPHO1-A novel phosphatase specifically expressed at sites of mineralisation in bone and cartilage. Bone 34:629–37. DOI: 10.1016/j.bone.2003.12.023.

Houston, DA, Myers, K, MacRae, VE, et al. (2016) The Expression of PHOSPHO1, nSMase2 and TNAP is Coordinately Regulated by Continuous PTH Exposure in Mineralising Osteoblast Cultures. Calcif Tissue Int 99:510–524. DOI: 10.1007/s00223-016-0176-9.

Hruska, KA, Mathew, S, Lund, R, et al. (2008) Hyperphosphatemia of chronic kidney disease. Kidney Int 74:148–57. DOI: 10.1038/ki.2008.130.

Huesa, C, Houston, D, Kiffer-Moreira, T, et al. (2015) The Functional co-operativity of Tissue-Nonspecific Alkaline Phosphatase (TNAP) and PHOSPHO1 during initiation of Skeletal Mineralization. Biochem Biophys Rep 4:196–201. DOI: 10.1016/j.bbrep.2015.09.013.

Jia, T, Olauson, H, Lindberg, K, et al. (2013) A novel model of adenine-induced tubulointerstitial nephropathy in mice. BMC Nephrol 14:116. DOI: 10.1186/1471-2369-14-116.

Kaleta, B (2019) The role of osteopontin in kidney diseases. Inflamm Res 68:93–102. DOI: 10.1007/s00011-018-1200-5.

Kazama, JJ, Iwasaki, Y & Fukagawa, M (2013) Uremic osteoporosis. Kidney Int Suppl (2011) :446–450. DOI: 10.1038/kisup.2013.93.

Khoshniat, S, Bourgine, A, Julien, M, et al. (2011) The emergence of phosphate as a specific signaling molecule in bone and other cell types in mammals. Cell Mol Life Sci 68:205–18. DOI: 10.1007/s00018-010-0527-z.

Lau, WL, Linnes, M, Chu, EY, et al. (2013a) High phosphate feeding promotes mineral and bone abnormalities in mice with chronic kidney disease. Nephrology Dialysis Transplantation 28:62–69. DOI: 10.1093/ndt/gfs333.

Lau, WL, Linnes, M, Chu, EY, et al. (2013b) High phosphate feeding promotes mineral and bone abnormalities in mice with chronic kidney disease. Nephrol Dial Transplant 28:62–9. DOI: 10.1093/ndt/gfs333.

Ma, YL, Cain, RL, Halladay, DL, et al. (2001) Catabolic effects of continuous human PTH (1--38) in vivo is associated with sustained stimulation of RANKL and inhibition of osteoprotegerin and gene-associated bone formation. Endocrinology 142:4047–54. DOI: 10.1210/endo.142.9.8356.

Metzger, CE, Swallow, E. & Allen, MR (2020) Elevations in Cortical Porosity Occur Prior to Significant Rise in Serum Parathyroid Hormone in Young Female Mice with Adenine-Induced CKD. Calcif Tissue Int 106:392–400. DOI: 10.1007/s00223-019-00642-w.

Metzger, CE, Swallow, EA, Stacy, AJ, et al. (2021) Adenine-induced chronic kidney disease induces a similar skeletal phenotype in male and female C57BL/6 mice with more severe deficits in cortical bone properties of male mice. PLoS One 16:e0250438. DOI: 10.1371/journal.pone.0250438.

Miller, MA, Chin, J, Miller, SC, et al. (1998) Disparate effects of mild, moderate, and severe secondary hyperparathyroidism on cancellous and cortical bone in rats with chronic renal insufficiency. Bone 23:257–266. DOI: https://doi.org/10.1016/S8756-3282(98)00098-2.

Mirza, MA, Hansen, T, Johansson, L, et al. (2009) Relationship between circulating FGF23 and total body atherosclerosis in the community. Nephrol Dial Transplant 24:3125–31. DOI: 10.1093/ndt/gfp205.

Moe, S, Drüeke, T, Cunningham, J, et al. (2006) Definition, evaluation, and classification of renal osteodystrophy: a position statement from Kidney Disease: Improving Global Outcomes (KDIGO). Kidney Int 69:1945–53. DOI: 10.1038/sj.ki.5000414.

Murali, SK, Andrukhova, O, Clinkenbeard, EL, et al. (2016a) Excessive Osteocytic Fgf23 Secretion Contributes to Pyrophosphate Accumulation and Mineralization Defect in Hyp Mice. PLoS Biol 14:e1002427. DOI: 10.1371/journal.pbio.1002427.

Murali, SK, Roschger, P, Zeitz, U, et al. (2016b) FGF23 Regulates Bone Mineralization in a 1,25(OH)2 D3 and Klotho-Independent Manner. J Bone Miner Res 31:129–42. DOI: 10.1002/jbmr.2606.

Nickolas, TL, Stein, EM, Dworakowski, E, et al. (2013) Rapid cortical bone loss in patients with chronic kidney disease. Journal of Bone and Mineral Research 28:1811–1820. DOI: https://doi.org/10.1002/jbmr.1916.

Ogirima, T, Tano, K, Kanehara, M, et al. (2006) Sex difference of adenine effects in rats: renal function, bone mineral density and sex steroidogenesis. Endocr J 53:407–13. DOI: 10.1507/endocrj.k05-009.

Qin, L, Raggatt, L. & Partridge, NC (2004) Parathyroid hormone: a double-edged sword for bone metabolism. Trends Endocrinol Metab 15:60–5. DOI: 10.1016/j.tem.2004.01.006.

Rendenbach, C, Yorgan, TA, Heckt, T, et al. (2014) Effects of extracellular phosphate on gene expression in murine osteoblasts. Calcif Tissue Int 94:474–83. DOI: 10.1007/s00223-013-9831-6.

Roberts, S, Narisawa, S, Harmey, D, et al. (2007) Functional involvement of PHOSPHO1 in matrix vesicle-mediated skeletal mineralization. J Bone Miner Res 22:617–27. DOI: 10.1359/jbmr.070108.

Shalhoub, V, Ward, SC, Sun, B, et al. (2011) Fibroblast growth factor 23 (FGF23) and alpha-klotho stimulate osteoblastic MC3T3.E1 cell proliferation and inhibit mineralization. Calcif Tissue Int 89:140–50. DOI: 10.1007/s00223-011-9501-5.

Shimada, T, Kakitani, M, Yamazaki, Y, et al. (2004) Targeted ablation of Fgf23 demonstrates an essential physiological role of FGF23 in phosphate and vitamin D metabolism. J Clin Invest 113:561–568. DOI: 10.1172/JCI19081.

Stenzel, I, Ziethe, K, Schurath, J, et al. (2003) Differential expression of the LePS2 phosphatase gene family in response to phosphate availability, pathogen infection and during development. Physiol Plant 118:138–146. DOI: 10.1034/j.1399-3054.2003.00091.x.

Tamura, M, Aizawa, R, Hori, M, et al. (2009) Progressive renal dysfunction and macrophage infiltration in interstitial fibrosis in an adenine-induced tubulointerstitial nephritis mouse model. Histochemistry and Cell Biology 131:483–490. DOI: 10.1007/s00418-009-0557-5.

Viau, A, El Karoui, K, Laouari, D, et al. (2010) Lipocalin 2 is essential for chronic kidney disease progression in mice and humans. J Clin Invest 120:4065–76. DOI: 10.1172/jci42004.

Whyte, MP (2008) Chapter 73-Hypophosphatasia: Nature ‘s Window on Alkaline Phosphatase Function in Humans, In: Bilezikian, JP, et al. (Eds), Principles of Bone Biology (Third Edition), San Diego, Academic Press. 1573–1598.

Yadav, MC, Simão, AMS, Narisawa, S, et al. (2011) Loss of skeletal mineralization by the simultaneous ablation of PHOSPHO1 and alkaline phosphatase function: A unified model of the mechanisms of initiation of skeletal calcification. Journal of Bone and Mineral Research 26:286–297. DOI: https://doi.org/10.1002/jbmr.195.

Yoshiko, Y, Candeliere, GA, Maeda, N, et al. (2007) Osteoblast autonomous Pi regulation via Pit1 plays a role in bone mineralization. Mol Cell Biol 27:4465–74. DOI: 10.1128/mcb.00104-07.

Zheng, CM, Zheng, JQ, Wu, CC, et al. (2016) Bone loss in chronic kidney disease: Quantity or quality? Bone 87:57–70. DOI: 10.1016/j.bone.2016.03.017.

