## Supplementary figures and images for "Increased PHOSPHO1 expression mediates cortical bone mineral density in renal osteodystrophy"

### Supplemental Fig S1

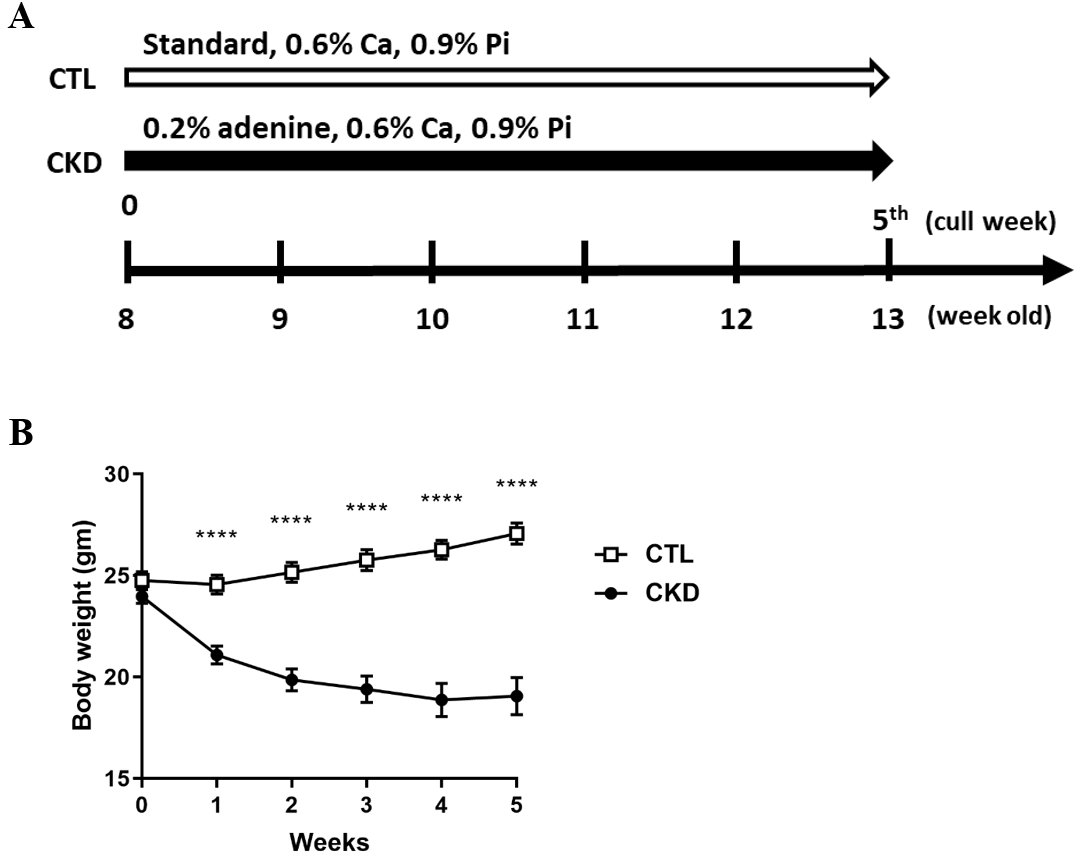

### Supplemental Fig S2

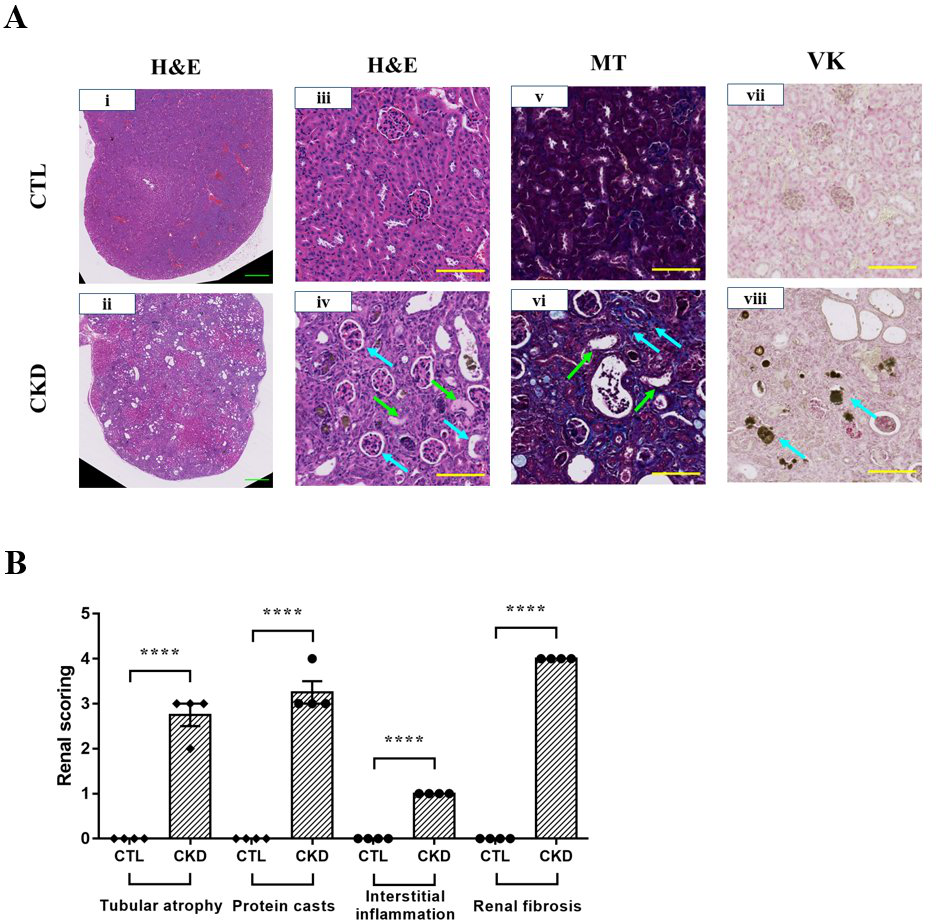

### Supplemental Fig S3

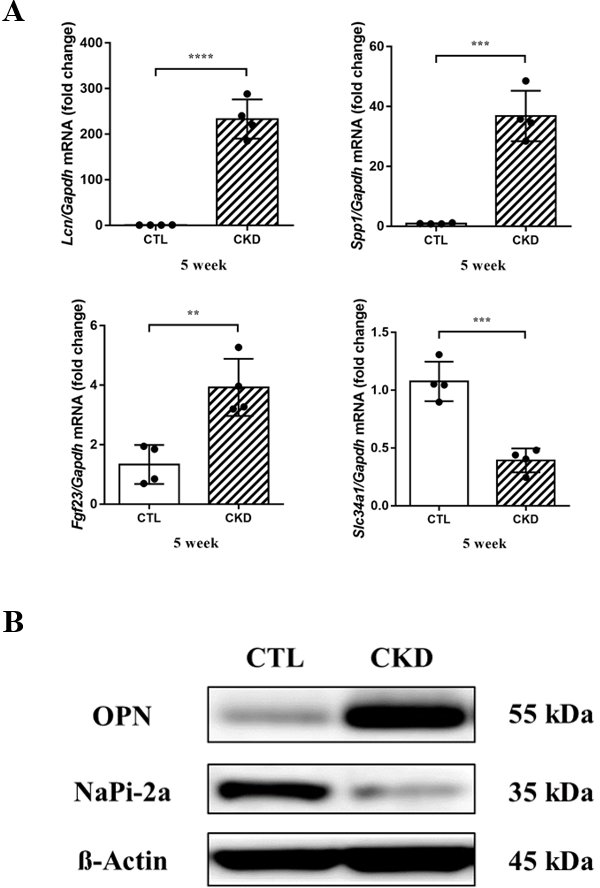

### Supplemental Fig S4

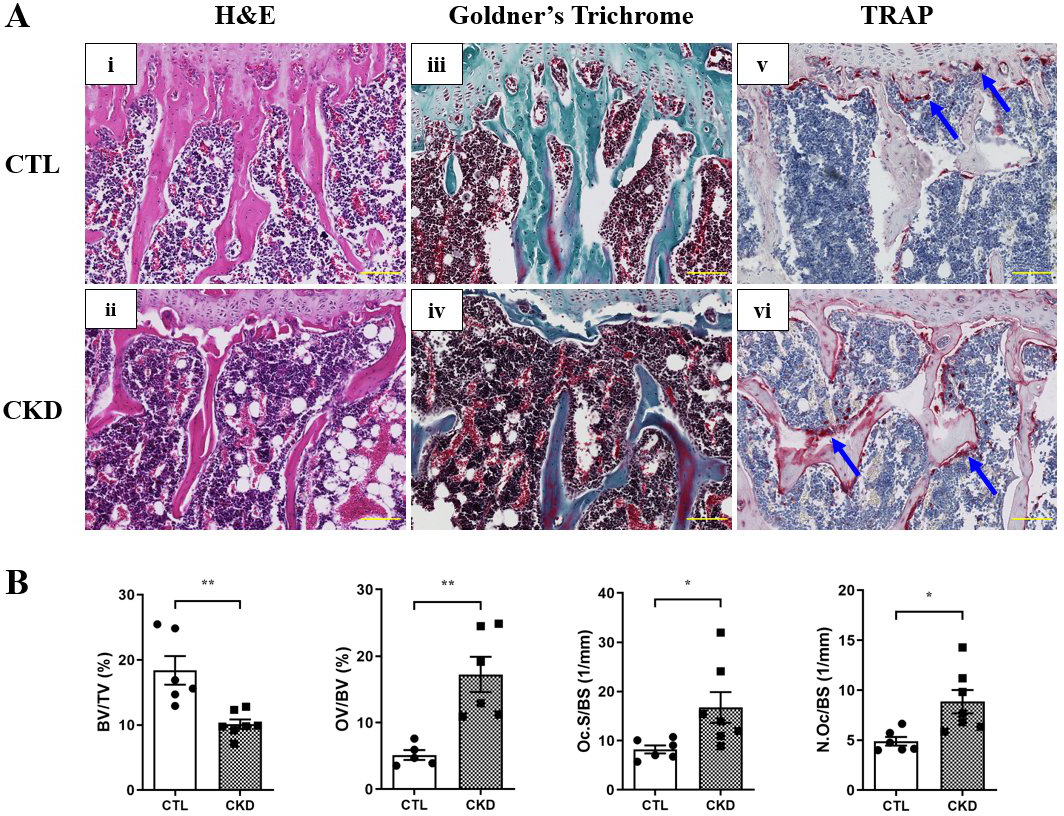

### Supplemental Fig S5

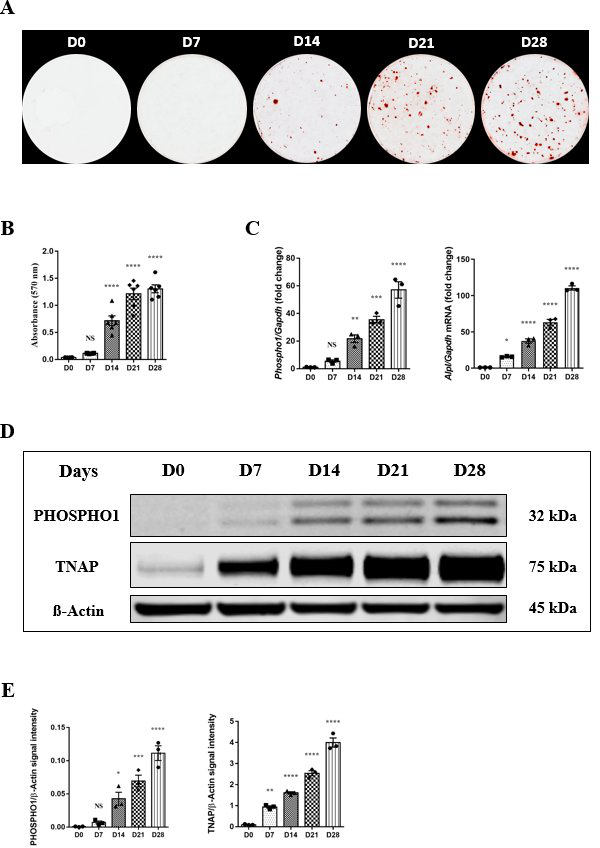

### Supplemental Fig S6

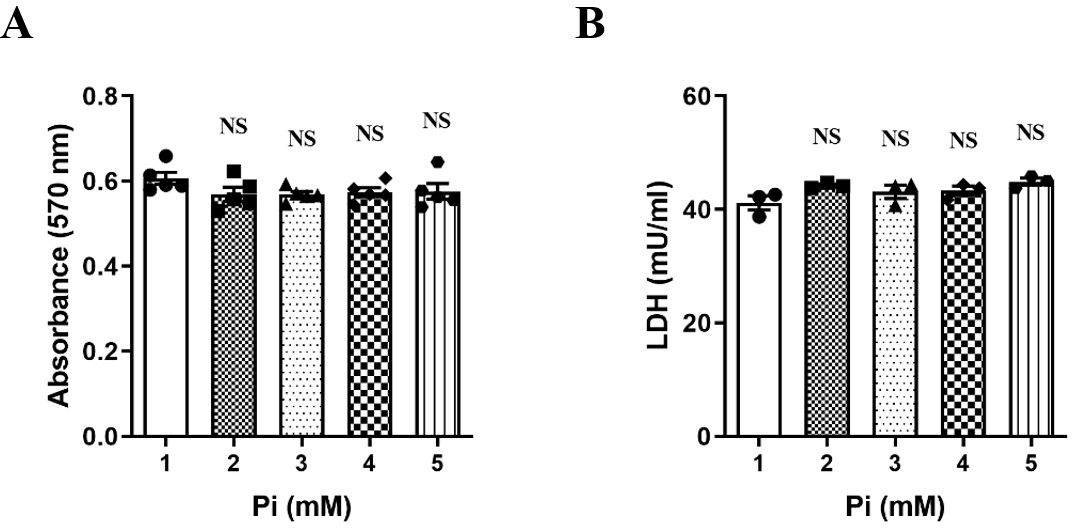
